# SpectralTAD: an R package for defining a hierarchy of Topologically Associated Domains using spectral clustering

**DOI:** 10.1101/549170

**Authors:** Kellen G. Cresswell, John C. Stansfield, Mikhail G. Dozmorov

## Abstract

The three-dimensional (3D) structure of the genome plays a crucial role in regulating gene expression. Chromatin conformation capture technologies (Hi-C) have revealed that the genome is organized in a hierarchy of topologically associated domains (TADs), sub-TADs, and chromatin loops. Identifying such hierarchical structures is a critical step in understanding regulatory interactions within the genome. Existing tools for TAD calling frequently require tunable parameters, are sensitive to biases such as sequencing depth, resolution, and sparsity of Hi-C data, and are computationally inefficient. Furthermore, the choice of TAD callers within the R/Bioconductor ecosystem is limited. To address these challenges, we frame the problem of TAD detection in a spectral clustering framework. Our `SpectralTAD` R package has automatic parameter selection, is robust to sequencing depth, resolution and sparsity of Hi-C data, and detects hierarchical, biologically relevant TAD structure. Using simulated and experimental Hi-C data, we show that `SpectralTAD` outperforms four state-of-the-art TAD callers. We demonstrate that TAD boundaries shared among multiple levels of the hierarchy were more enriched in classical boundary marks, such as CTCF, RAD21, and more conserved across cell lines and tissues. In contrast, boundaries of primary TADs, defined as TADs which cannot be split into sub-TADs, showed less enrichment and conservation, suggesting their more dynamic role in genome regulation. In summary, we present a simple, fast, and user-friendly R package for robust detection of TAD hierarchies supported by biological evidence. `SpectralTAD` is available on Bioconductor, http://bioconductor.org/packages/SpectralTAD/.

## Introduction

The introduction of chromatin conformation capture technology and its high-throughput derivative Hi-C enabled researchers to accurately model chromatin interactions across the genome and uncover the non-random 3D structures formed by folded genomic DNA [1–3]. The structure and interactions of the DNA in 3D space inside the nucleus has been shown to shape cell type-specific gene expression [3–5], replication [6], guide X chromosome inactivation [7], and regulate the expression of tumor suppressors and oncogenes [8–10].

Topologically Associated Domains (TADs) refer to a common structure uncovered by Hi-C technology, characterized by groups of genomic loci that have high levels of interaction within the group and minimal levels of interaction outside of the group [1,7,11,12]. TAD boundaries were found to be enriched in CTCF (considering the directionality of its binding) and other architectural proteins of cohesin and mediator complex (e.g., STAG2, SMC3, SMC1A, RAD21, MED12) [13–15], marks of transcriptionally active chromatin (e.g., DNAse hypersensitive sites, H3K4me3, H3K27ac, H3K36me3 histone modifications) [16, 17], actively transcribed and housekeeping genes [18]. From a regulatory perspective, TADs can be thought of as isolated structures that serve to confine genomic activity within their walls, and restrict activity across their walls. This confinement has been described as creating “autonomous gene-domains,” essentially partitioning the genome into discrete functional regions [17, 19].

TADs organize themselves into hierarchical sets of domains [20–23]. These hierarchies are characterized by large “meta-TADs” that contain smaller sub-TADs and chromatin loops. To date, most methods were developed to find these single meta-TADs instead of focusing on the hierarchy of the TAD structures [24–26]. While interesting insights can be gleaned from the meta-TADs, work has shown that smaller sub-TADs are specifically associated with gene regulation [19,27,28]. For example, it has been found that genes associated with limb malformation in rats are specifically controlled through interactions within sub-TADs [28]. These results highlight the importance of identifying the full hierarchy of TADs.

Several methods have been designed to call hierarchical TADs (Supplemental Material). However, most algorithms require tunable parameters [25,29,30] that, if set incorrectly, can lead to a wide variety of results. Many tools have been shown to highly depend on sequencing depth and chromosome length (reviewed in [31]). Furthermore, the time complexity of many algorithms is often prohibitive for detecting TADs on a genome-wide scale. Also, many tools are not user-friendly and lack clear documentation [30, 32], with some methods even lacking publicly available code [21]. Furthermore, the choice of TAD callers in R/Bioconductor ecosystem remains limited (Supplemental Material).

Our goal was to develop a simple data-driven method to detect TADs and uncover hierarchical sub-structures within these TADs. We propose a novel method that exploits the graph-like structure of the chromatin contact matrix and extend it to find the full hierarchy of sub-TADs, limited only by the resolution of Hi-C data. The method employs a modified version of the multiclass spectral clustering algorithm [33], and uses a sliding window based on the commonly used 2Mb biologically maximum TAD size [11, 34]. We introduce a novel method for automatically choosing the number of clusters (TADs) based on maximizing the average silhouette score [35]. We show that this approach finds TAD boundaries with more significant enrichment of known boundary marks than those called by other TAD callers. We then extend the method to find hierarchies of TADs and demonstrate their biological relevance. Our method provides a parameterless approach, efficiently operating on matrices in text format with consistent results regardless of the level of noise, sparsity, and resolution of Hi-C data. The method is fast and scales linearly with the increasing amount of data. Our method is implemented in the `SpectralTAD` R package, freely available on GitHub (https://github.com/dozmorovlab/SpectralTAD) and Bioconductor (http://bioconductor.org/packages/SpectralTAD/).

## Results

### An overview of the SpectralTAD algorithm

SpectralTAD takes advantage of the natural graph-like structure of Hi-C data, allowing us to treat the Hi-C contact matrix as an adjacency matrix of a weighted graph. This interpretation allows us to use a spectral clustering-based approach, modified to use gaps between consecutive eigenvectors as a metric for defining TAD boundaries (Methods). We implement a sliding window approach that increases the stability of spectral clustering and reduces computation time. This approach detects the best number and quality of TADs in a data-driven manner by maximizing the number of internal contacts within TADs and minimizing those between TADs. To achieve this, we maximize a clustering metric called silhouette score that measures within TAD similarity and penalizes for high similarity between TADs.

### Defining hierarchical TADs and boundaries

We distinguish hierarchical types of TADs by their position with respect to other TADs. Primary TADs, or “meta-TADs,” are defined as the top-level TADs that are not enclosed within other TADs (Figure 1A). Conversely, we define “sub-TADs” as TADs detected within other TADs. We further refine the definition of sub-TADs to describe the level of hierarchy in which a sub-TAD is contained. Secondary TADs refer to sub-TADs which are contained within a primary TAD; tertiary TADs correspond to sub-TADs that are contained within two TADs and so on (Figure 1A). Unless specified otherwise, we report results concerning primary TADs.

**Figure 1.**
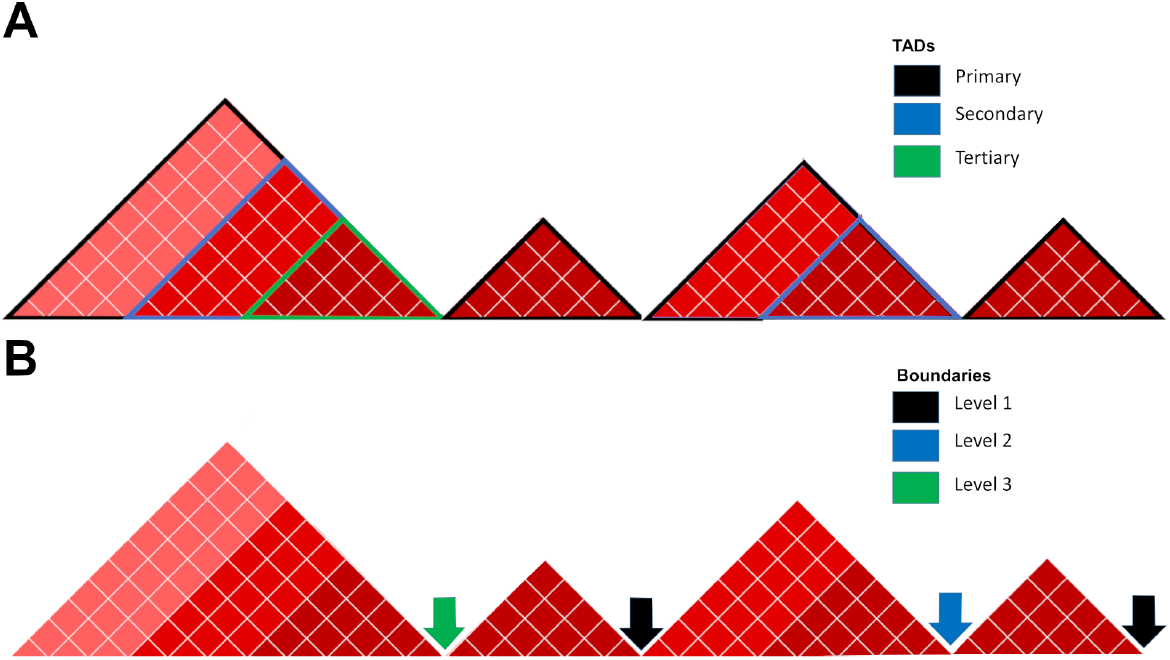
Hierarchy of TADs and boundaries. A) TADs not enclosed within other TADs are defined as primary, while TADs contained within other TADs are defined as secondary, tertiary, etc. B) Boundaries are defined as the rightmost point of a given TAD. Boundaries belonging to a single TADs are defined as level 1, while boundaries shared by two, three, TADs are defined as level 2, 3, etc.

TAD boundaries represent another important element to be considered within the hierarchy. Using the terminology introduced in An et al. [36], we define a level 1 boundary as a TAD boundary belonging to a single TAD, irrespective of the TAD type. Level 2 and level 3 boundaries correspond to boundaries that are shared by two or three TADs, respectively (Figure 1B).

An additional type of region is a gap, which refers to an area where there are no TADs present either due to a lack of sequencing depth, a centromere, or simply a lack of organization (Supplemental Methods). The percentage of non-centromeric gaps varies across chromosomes and resolutions (Supplemental Table S1), being 19.9% on average. In general, we observe that data at higher resolution (e.g., 10kb) have the highest percentages of gaps due to sparsity. In our analysis, TADs are allowed to span the non-centromeric gaps.

### Comparison of TAD quality

A TAD boundary detection method (“TAD detection” hereafter) must be robust to sparsity and noise in Hi-C data, detect consistent TADs across sequencing depths and resolutions, and the TADs must be biologically and statistically meaningful. To compare the concordance of TAD boundaries identified by different TAD callers under different conditions, we used the Jaccard similarity metric. For comparing TAD boundaries identified at different resolutions, we used a modified Jaccard similarity metric (Supplemental Figure S1, Supplemental Material). Using simulated and experimental Hi-C data, we compared our method, `SpectralTAD`, with two single-level R-based TAD callers (`TopDom` and `HiCSeg`), a hierarchical R-based TAD caller (`rGMAP`) and a Python-based hierarchical TAD caller (`OnTAD`).

An important property of TAD detection methods is the ability to detect a hierarchy of TAD structures [3,12,21,22,27,37]. Among R packages, `rGMAP` allows for the detection of two levels of TAD hierarchy. Our method, `SpectralTAD`, can detect deeper levels of hierarchy, though we limit it to three in the current paper (Supplemental Figure S2). Similarly to `SpectralTAD OnTAD`, Using simulated and experimental data, we compared the robustness of hierarchical TAD detection and defined properties of hierarchical TAD boundaries.

Multiple studies have demonstrated an enrichment of various genomic annotations at TAD boundaries [11,17,21,24]. We quantified the biological relevance of TAD boundaries by using a permutation test to determine their enrichment in transcription factor binding sites, histone modification marks and chromatin segmentation states (Supplemental Methods, Supplemental Table S2).

### ICE-normalized and raw Hi-C data are better suited for TAD detection

Sequence- and technology-driven biases may be present in Hi-C matrices [38–41]. Consequently, numerous normalization methods have been developed [2,40–45]. However, their effect on the quality of TAD detection has not been explored.

We investigated the effect of four normalization methods, Knight-Ruiz (KR), iterative correction and eigenvector decomposition (ICE), and square root vanilla coverage (sqrtVC) on TAD detection using `SpectralTAD`. Using simulated matrices with the ground-truth TADs, we found that all normalization methods marginally degraded the performance of `SpectralTAD` under different levels of noise, sparsity and downsampling (Figure 2A-C). These results suggest that the use of raw Hi-C data is appropriate for the detection of TAD boundaries.

**Figure 2.**
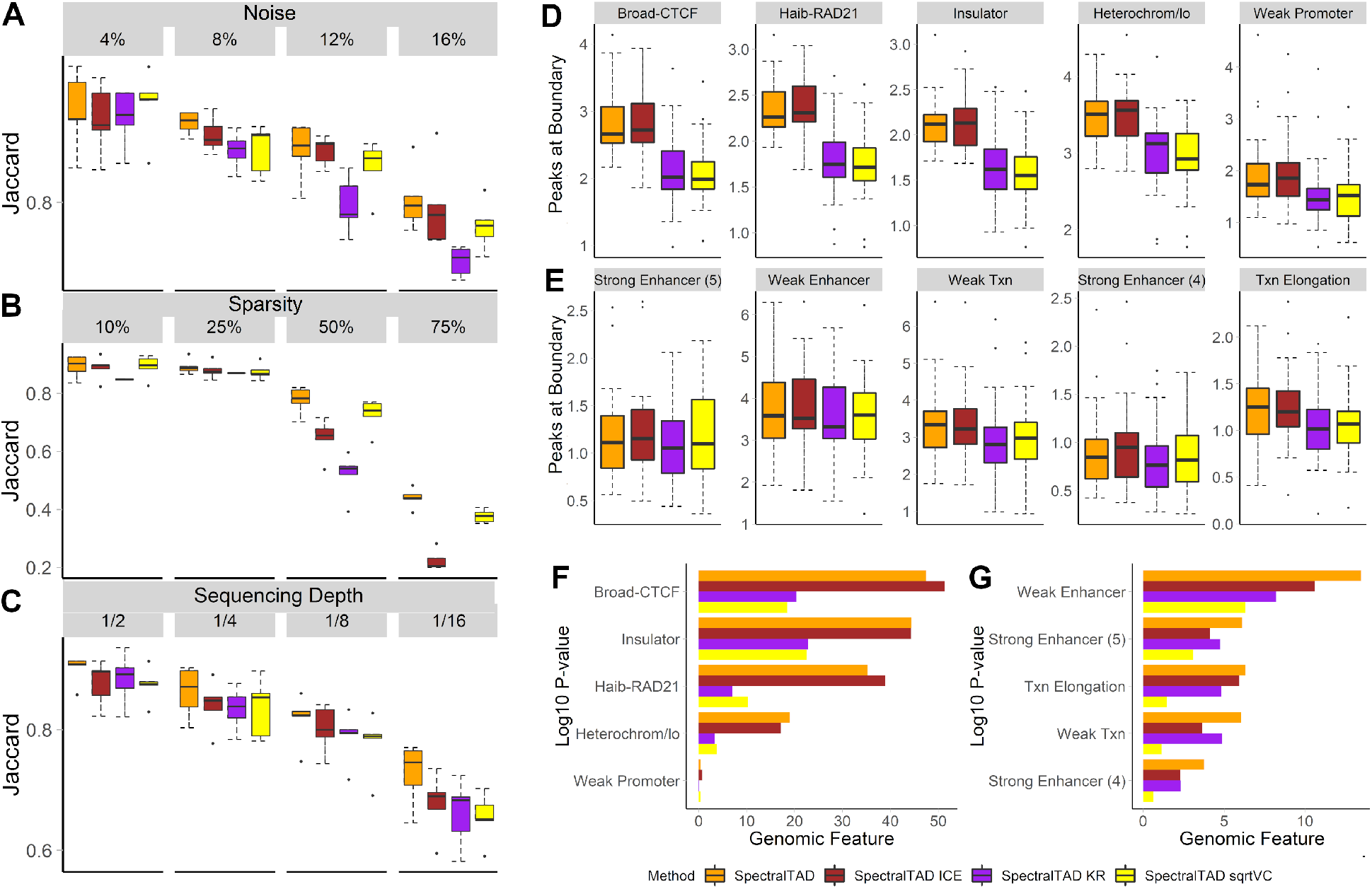
The effect of normalization on TAD consistency and enrichment. To test for robustness to noise, sparsity, and sequencing depth, simulated Hi-C matrices were used as-is, KR-, ICE- and square root VC-normalized. TADs detected by `SpectralTAD` were compared with the ground-truth TADs using the Jaccard similarity metric. The effect of normalization was assessed at different levels of noise (A, the percentage of the original contact matrix modified by adding a constant), sparsity (B, the percentage of the original contact matrix replaced with zero), and downsampling (C, the fraction of contacts kept, see Methods). Using the raw and normalized data from Gm12878 cell line at 25kb resolution, enrichment of genomic annotations within 50kb regions flanking a TAD boundary on both sides were assessed using a permutation test. The average number of annotations for enriched (D) and depleted (E) genomic features and the permutation p-values corresponding to enrichment (F), and depletion (G) for the top five most enriched/depleted genomic annotations are shown. Results averaged across chromosome 1-22 are shown.

Using the experimental Hi-C data from Gm12878 cell line, we found that ICE normalization only marginally affected the average number and width of TADs, and these results were consistent across resolutions (Supplemental Figure S3A). In contrast, KR, and sqrtVC normalization resulted in a larger variability in TAD widths across chromosomes and between resolutions (Supplemental Figure S3B). We also assessed the average number and the enrichment (permutation test) of genomic annotations at TAD boundaries detected from unnormalized, KR-, ICE-, and sqrtVC-normalized data. The average number of genomic annotations was not significantly different in TAD boundaries detected from raw and ICE-normalized data as compared with those from KR- and sqrtVC-normalized, where the number of annotations was significantly less (Figure 2D). We found CTCF, RAD21, “Insulator,” and “Heterochromatin” states to be significantly enriched in TAD boundaries, and this enrichment was frequently more significant in TAD boundaries detected from the ICE-normalized data (Figure 2F). Similarly, “enhancer”-like chromatin states were significantly depleted at TAD boundaries, and this depletion was more pronounced in boundaries detected from raw data (Figure 2G). The enrichment results were consistent across resolutions (Supplemental Figure S3C-F). These results suggest that both ICE-normalized and raw Hi-C data are suitable for the robust detection of biologically relevant TADs. Based on these results, and the fact that previous studies showed graph-based TAD identification methods work well un-normalized HiC data [46, 47], consequent results are presented with the use of raw Hi-C data.

### SpectralTAD identifies more consistent TADs than other methods

Using simulated matrices, we compared the performance of `SpectralTAD` with `rGMAP, TopDom, OnTAD` and `HiCSeg` at different noise levels. We found that both `SpectralTAD` and `TopDom` had a significantly higher agreement with the ground truth TADs than `rGMAP` across the range of noise levels (Figure 3A). To better understand the poor performance of `rGMAP`, we hypothesized that inconsistencies might arise due to the “off-by-one” errors that occur when, by chance, a TAD boundary may be detected adjacent to the true boundary location. We analyzed the same data using TAD boundaries flanked by 50kb regions. Expectedly, the performance of all TAD callers, including `rGMAP`, increased; yet, the performance of `rGMAP` remained significantly low (Supplemental Figure S4A). At low level of noise, `HiCseg` detected highly consistent TAD boundaries; however, these TAD boundaries were least biologically relevant, detailed below. In summary, these results suggest that, with the presence of high noise levels, a situation frequent in experimental Hi-C data, `SpectralTAD` performs better than other TAD callers in detecting true TAD boundaries.

**Figure 3.**
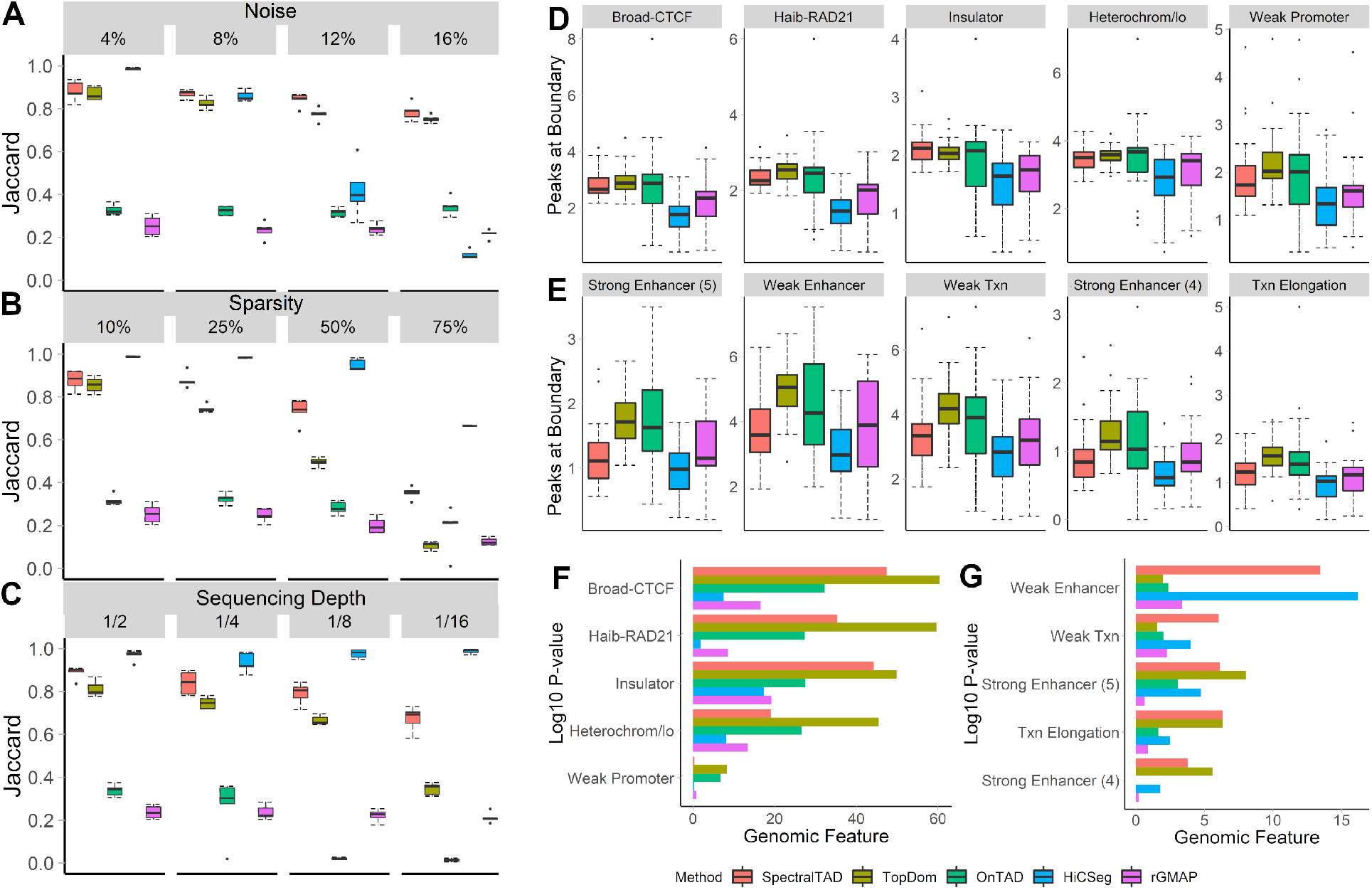
The comparison of SpectralTAD and other TAD callers regarding TAD consistency and biological significance. To test for robustness to noise, sparsity, and sequencing depth, TADs were called from simulated Hi-C matrices using `SpectralTAD` and four other TAD callers. They were compared with the ground-truth TADs using the Jaccard similarity metric. The performance was assessed at different levels of noise (A), sparsity (B), and downsampling (C, see Methods). Using the raw data from Gm12878 at 25kb resolution, enrichment of genomic annotations within 50kb regions flanking a TAD boundary on both sides was assessed using a permutation test. The average number of annotations for enriched (D) and depleted (E) genomic features and the permutation p-values corresponding to enrichment (F), and depletion (G) for the top five most enriched/depleted genomic annotations are shown. Results averaged across chromosome 1-22 are shown.

We similarly investigated the effect of sparsity on the performance of the TAD callers. Expectedly, the average Jaccard similarity decreased for all TAD callers with the increased level of sparsity (Figure 3B). `SpectralTAD` outperformed all TAD callers except `HiCseg` at all sparsity levels. We further tested whether accounting for the “off-by-one” error improves the performance; the performance of `SpectralTAD` remained superior (Supplemental Figure S4B). These results demonstrate the robustness of `SpectralTAD` to sparsity.

TAD callers should be robust to changes in sequencing depth. We introduced four levels of downsampling into simulated matrices and compared the detected TADs with the ground truth TADs. Downsampling involves removing contacts at random, simulating sequencing depth. Expectedly, the average Jaccard similarity degraded for all TAD callers with the increased level of downsampling (Figure 3C). Notably, the performance of `SpectralTAD` was consistently higher than that of for other TAD callers except `HiCseg`. Similar observations were true when accounting for the “off-by-one” error (Supplemental Figure S4C). Despite seemingly good performance of `HiCseg`, the biological relevance of TAD boundaries it detects is low, as shown below (Figure 3). These observations, along with the results concerning sparsity and noise, suggest that with realistic levels of variation and noise in Hi-C data the performance of `SpectralTAD` is better than other TAD callers.

### SpectralTAD outperforms other TAD callers in finding biologically relevant TAD boundaries

To evaluate the biological relevance of TAD boundaries detected by `SpectralTAD` and the other TAD callers, we evaluated their enrichment in genomic annotations known to be associated with TAD boundaries. We found that the TAD boundaries called by `SpectralTAD, TopDom`, and `OnTAD` had a significantly higher number of CTCF and RAD21, “Insulator,” and “Heterochromatin” annotations than those called by `HiCseg` and `rGMAP` (Figure 3D). Consequently, these marks were more enriched at TAD boundaries detected by `SpectralTAD` and `TopDom` as compared with the other TAD callers (Figure 3F). In terms of depleted genomic annotations, “enhancer”-like chromatin states were underrepresented at TAD boundaries, and this depletion was highly significant for boundaries detected by `SpectralTAD` (Figure 3E, G). Notably, the TAD boundaries detected by `HiCseg` had the lowest number of these genomic annotations. They also exhibited the lowest level of enrichment and depletion (Figure 3E, G). These results suggest that, despite robustness to noise, sparsity, and sequencing depth, HiCseq detects boundaries that are less biologically relevant in terms of known TAD biology. The performance of `SpectralTAD` and other callers was consistent at different resolutions (Supplemental Figure S4D-G, Supplemental Table S3). In summary, these results suggest that `SpectralTAD` outperforms other TAD callers in detecting biologically relevant TAD boundaries.

### SpectralTAD identifies consistent TADs across resolutions of Hi-C data

It is imperative that TAD boundaries called at different resolutions of Hi-C data are consistent, or one risks finding vastly different TAD boundaries despite the data being the same. Using the Gm12878 Hi-C data at 10kb, 25kb, and 50kb resolutions, we estimated the average number and width of TADs called by `SpectralTAD, TopDom, HiCseg, OnTAD`, and `rGMAP`. As resolution of Hi-C data increased, the average number of TADs decreased for all but `SpectralTAD` (Supplemental Figure S5A). Similarly, the average width of TADs increased for all but `SpectralTAD` TAD callers (Supplemental Figure S5B). We further compared the consistency of TADs detected in 50kb vs. 25kb, 50kb vs. 10kb, 25kb vs. 10kb resolution comparisons. We found that, for nearly all comparisons, `SpectralTAD` and `HiCseg` had significantly higher consistency quantified by modified Jaccard statistics than the other TAD callers (Supplemental Figure S5C). These results show that `SpectralTAD` identifies consistent TADs at different resolutions of Hi-C data.

### The hierarchical structure of TADs is associated with biological relevance

Having established the strong performance of `SpectralTAD`, we investigated the biological importance of the hierarchy of TAD boundaries detected by it. We tested the relationship between the number of times a TAD boundary occurs in a hierarchy (Figure 1B) and enrichment of genomic annotations. We hypothesized that TAD boundaries shared by two or more TADs (Level 2 and 3 boundaries) would be more biologically important, hence, harbor a larger number of key markers such as CTCF and RAD21. We found that this is indeed the case, as illustrated by a significant increase in the average number of CTCF and RAD21 annotations, and “Insulator,” and “Heterochromatin” states around Level 2 and 3 boundaries as compared with Level 1 boundaries (Figure 4A). A similar trend was observed in the stronger enrichment, of Level 2 and 3 TAD boundaries in those annotations (Figure 4C). TAD boundaries at all levels of the hierarchy were similarly depleted in the “enhancer”-like annotations (Figure 4B, D), although these depletions were more significant for the Level 3 TAD boundaries. These observations were consistent across resolutions (Supplemental Figure S6). Our results agree with previous research that has shown a positive correlation between the number of sub-TADs sharing a boundary and the number of biologically relevant genomic annotations at that boundary [36, 48] and confirm that `SpectralTAD` identifies a biologically relevant hierarchy of TAD boundaries.

**Figure 4.**
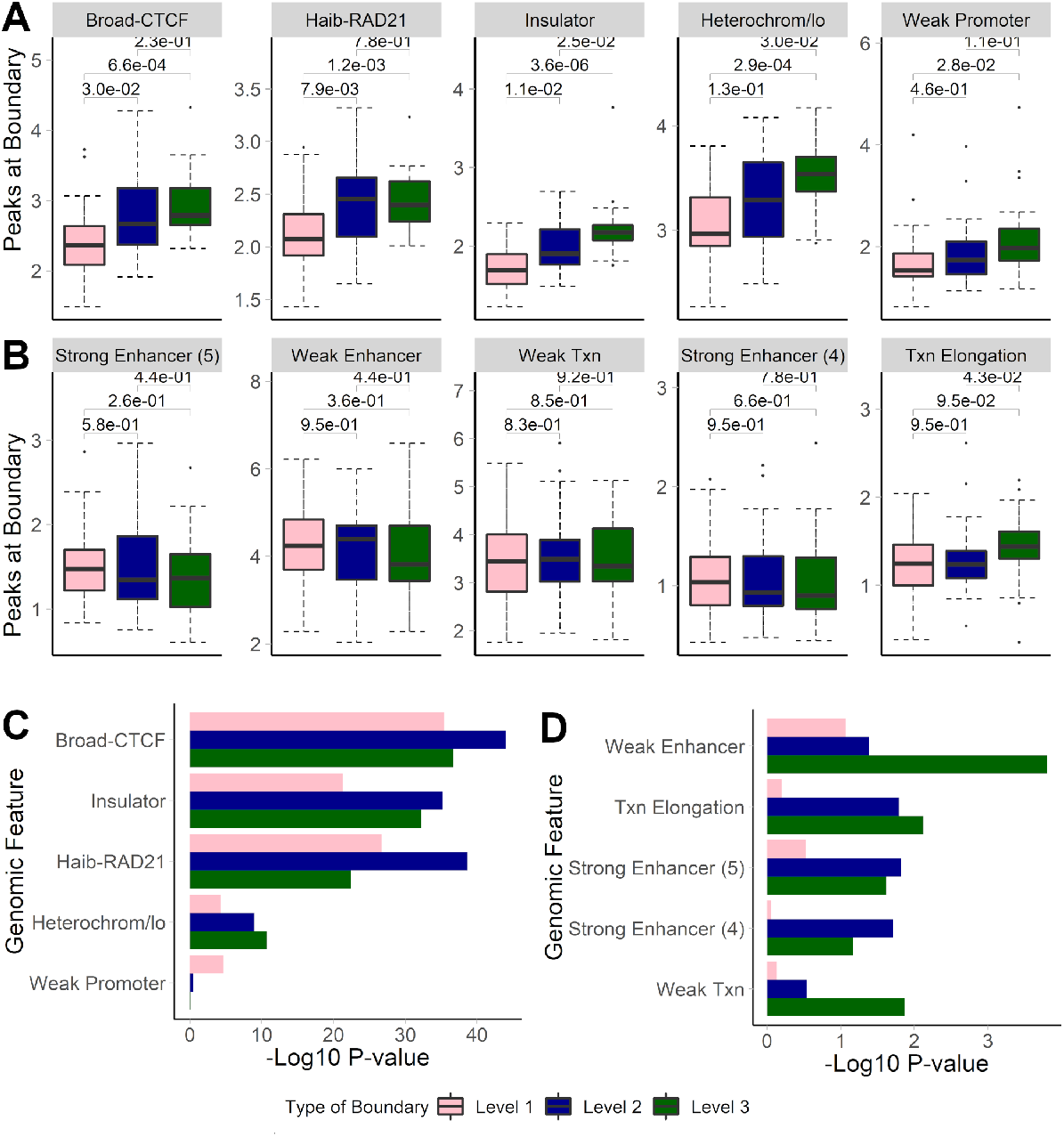
The effect of the hierarchy of TAD boundaries detected by SpectralTAD on the average number of annotations in enriched (A) & depleted (B) genomic markers and on enrichment (C) and depletion (D) for different genomic annotations. Results for TAD boundaries detected as level 1, 2, and 3 boundaries are shown. Genomic annotations were considered within 50kb regions flanking a boundary on both sides. Wilcoxon test p-values, summarized over chromosomes, are shown in panel A & B and aggregated p-values, using the Fisher’s method, are shown for panels C & D. Raw data from the GM12878 cell line, chromosome 1-22, 25kb resolution.

### TADs identified by SpectralTAD are conserved across cell-line and tissues

Previous studies reported relatively high conservation of TAD boundaries identified in different tissues and cell types, with the reported Jaccard statistics ranging from 0.21 to 0.30 [3]. We compared TAD boundaries called by `SpectralTAD` across various tissues and cell types (Supplemental Table S4, [49]). The Jaccard for all TADs, ignoring hierarchy, between cell-line samples ranged from 0.33 to 0.73 with a mean of 0.45 (SD = 0.08). The Jaccard between tissues ranged from 0.21 to 0.38 with a mean of 0.27 (SD = 0.03), significantly lower than that of cell lines (Wilcoxon p-value < 0.0001). The lower conservation of TADs called from tissue samples is expected as cell lines come from a “pure” single source while tissues are a mixture of different cells. These results were summarized in heatmaps, comparing different cell lines (Supplemental Figure S7A) and tissues (Supplemental Figure S7B) according to Jaccard similarity. Hierarchical clustering of cell type-specific samples by the Jaccard similarity of their TADs, ignoring hierarchy, identified the expected associations between cell type-specific data, with replicates clustering together and cell types being distinct (Supplemental Figure S7A). To a lesser extent, these results were similar in tissue-specific samples (Supplemental Figure S7B). In summary, these results show conservation of TAD boundaries called by `SpectralTAD` across tissues and cell lines similar to previously reported results.

### Hierarchy of TADs and boundaries affect conservation of TADs

Following our definition of TADs (Primary, Secondary, and Tertiary, Figure 1A), we hypothesized that primary TADs would be better conserved than Secondary or Tertiary TADs. The primary TADs are detected during the first pass of the algorithm; hence, they are robustly supported by the underlying data and expected to reproduce across different datasets. Indeed, the average Jaccard for Primary, Secondary, and Tertiary TADs across cell types was 0.42, 0.40, and 0.35, respectively (Supplemental Table S5), and this decrease was significant (Wilcoxon p-value < 0.0001). These observations were consistent when analyzing TADs called from tissue samples, although the average Jaccard coefficients for Primary, Secondary, and Tertiary TADs were significantly lower (Supplemental Table S5). These results demonstrate that Primary TADs are the most conserved across cell types and tissues.

We hypothesized that Level 3 TAD boundaries (Figure 1B, boundaries that are shared by three TADs), besides showing higher biological significance (Figure 4), will be better conserved. Indeed, the Jaccard coefficient of Level 3 TAD boundaries called in cell types was significantly higher (0.30) than that of Level 2 (0.23) and Level 1 (0.23) boundaries (Wilcoxon p-value ranging from 0.034 to <0.0001, Supplemental Table S5). These results were also observed in TAD boundaries called in tissue types. One possibility of the lower Jaccard coefficient for Level 1 and 2 boundaries is that they may change their assignment due to higher probability of detection of sub-TADs in different datasets. In summary, these results demonstrate that boundaries shared by several TADs have high biological significance and are better conserved across cell types and tissues.

### SpectralTAD is the fastest TAD caller for high-resolution data

We evaluated the runtime performance of `SpectralTAD, TopDom, OnTAD, rGMAP` and `HiCSeg. SpectralTAD` showed comparable performance with `TopDom` and was faster than `rGMAP` at all resolutions (Supplemental Figure S8A). Specifically, `SpectralTAD` takes ~45 seconds to run with 25kb data and ~4 minutes to run on 10kb data for the entire GM12878 genome. By comparison, `TopDom` takes ~1 minute to run on 25kb data but ~13 minutes on 10kb data. `OnTAD` takes ~4 minutes to run on 25kb data and ~30 minutes on 10kb data. `rGMAP` takes ~12 minutes on 25kb data and ~47 minutes on 10kb data. We find that HiCSeg is prohibitively slow, taking ~609 minutes on 25kb data and multiple days to run on 10kb data with chromosome 1 taking over 24 hours alone. Importantly, our method scales nearly linearly with the size of the data (see Methods), making it amenable for fast processing of data at higher resolutions. Furthermore, when parallelized, `SpectralTAD` is several orders of magnitude faster than other TAD callers (Supplemental Figure S8A), e.g., with the entire genome taking 1 second to run for 25kb data when using four cores. We demonstrate that our method has a linear complexity *O*(*n*) (Supplemental Methods) making it scalable for large Hi-C datasets. In summary, these results demonstrate that `SpectralTAD` is significantly faster than `TopDom, rGMAP` and `HiCSeg`, providing near-instant results when running on multiple cores.

## Discussion

We introduce the `SpectralTAD` R package implementing a spectral clustering-based approach that allows for fast TAD calling and scales well to high-dimensional data. The method was benchmarked against four TAD callers implemented in R - `TopDom` and `OnTAD` that detect single-level `TADs`, and `HiCseg` and `rGMAP` hierarchical TAD caller. We show better performance of `SpectralTAD` vs. the other TAD callers in nearly all conditions. We also demonstrate that `SpectralTAD` is more robust to sparsity, sequencing depth and resolution. We show that `SpectralTAD` can robustly detect hierarchical TAD boundaries. Furthermore, we demonstrate different levels of TAD hierarchy to be differentially associated with known marks of TAD boundaries, highlighting their distinct biological roles and the importance of TAD hierarchy in general. The clear superiority of `SpectralTAD` regarding running speed and robustness to data irregularities suggests its use as the new gold-standard of hierarchical TAD callers in the R ecosystem.

The performance of `SpectralTAD` was frequently better, but not always superior to that of `HiCseg`. The better performance of `HiCseg` under different levels of noise, sparsity, and sequencing depth in some cases may be explained by the fact that `HiCseg` identifies non-hierarchical TAD boundaries at once, thus detecting the maximum number of them in one run. `SpectralTAD`, on the other hand, defines a hierarchy of primary, secondary, etc., TADs, restricted to the first three levels in the current analysis. Thus, TADs at a deeper level may have been detected by `HiCseg` inflating its performance. However, TAD boundaries identified by `HiCseg` were significantly less associated with known marks of TAD boundaries, undermining their biological relevance. We suggest that the tradeoff between robustness and biological relevance of TAD boundaries should be made for the latter, with `SpectralTAD` providing the optimal balance.

One overarching limitation with non-hierarchical TAD callers like `TopDom` and `HiCSeg` is their inability to capture all TADs in a dataset. While methods like `TopDom` may find biologically relevant TADs, they cannot account for the common situation where sub-TADs occur within a TAD. In the case of TADs enclosing sub-TADs, non-hierarchical callers are forced to make a choice, often far from optimal (Supplemental Figure S2). We suggest that even when the hierarchy of TADs is not essential, hierarchical TAD callers like `SpectralTAD` should be used for maximally accurate reconstruction of TADs at first level of the hierarchy.

In summary, we show that `SpectralTAD` is a robust method for defining the hierarchy of TAD boundaries. This method improves upon previous work showing the potential of spectral clustering for finding structures in Hi-C data while introducing modifications to make these methods practical for users. Specifically, we introduce two novel modifications to spectral clustering, the eigenvector gap and windowing, which can be used to quickly and accurately find changes in the pattern for ordered data. By releasing `SpectralTAD` as an open source R package, we aim to provide a user-friendly and accurate tool for hierarchical TAD detection.

## Methods

### Data Sources

Experimental Hi-C matrices from the Gm12878 cell line ([3] at 50kb, 25kb, and 10kb) and 35 different cell line and tissue samples ([49], 40kb resolution) were downloaded from Gene Expression Omnibus (GEO, Supplemental Table S4). The Gm12878 data is the “primary+replicate” data from [3] and was generated by combining results from two experiments by counting total contacts and binning the genome, using increasing sizes, until 80% of bins contained over 1000 contacts. 25 simulated matrices with manually annotated TADs ([50], 40kb resolution) were downloaded from the `HiCToolsCompare` repository (Supplemental Table S4). Data for chromatin states, histone modification and transcription factor binding sites (TFBS) were downloaded from the UCSC genome browser database [51]. Given the fact that some transcription factors have been profiled by different institutions (e.g., CTCF-Broad, CTCF-Uw, and CTCF-Uta), we selected annotations most frequently enriched at TAD boundaries (typically, CTCF-Broad, RAD21-Haib). All genomic annotation data were downloaded in Browser Extensible Data (BED) format using the hg19/GRCh37 genome coordinate system (Supplemental Table S2).

### Windowed spectral clustering

#### Hi-C data representation

Chromosome-specific Hi-C data is typically represented by a chromatin interaction matrix *C* (referred hereafter as “contact matrix”) binned into regions of size *r* (the resolution of the data). Entry *C_ij_* of a contact matrix corresponds to the number of times region *i* interacts with region *j*. The matrix *C* is square and symmetric around the diagonal representing self-interacting regions. Our method relies on the fact that the 3D chromosome can be thought of as a naturally occurring graph. Traditionally, a graph *G*(*V, E*) is represented by a series of nodes *V* connected by edges *E*. These graphs are summarized in an adjacency matrix *A_i_j* where entry *ij* indicates the number of edges between node *i* and node *j*. We can think of the contact matrix as a naturally occurring adjacency matrix (i.e., *C_ij_* = *A_ij_*) where each genomic locus is a node and the edges are the number of contacts between these nodes. This interpretation of the contact matrix allows us to proceed with spectral clustering.

#### Sliding Window

To avoid performing spectral clustering on the entire matrix, which is highly computationally intensive, we apply the spectral clustering algorithm to submatrices defined by a sliding window across the diagonal of the entire matrix. The size of the window (the number of bins defining a submatrix) is based on the maximum possible TAD size of 2mb [11, 52]. In practice, the size of the window *w* is equal to 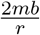 where *r* is the resolution of the data. For example, at the 10kb resolution, we would have a window size of 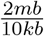 or simply 200 bins. Following the guidelines of previous works on the minimum TAD size, we set a minimum window size of 5 bins [11,53–55].

The restriction in window size means that the maximum resolution at which the algorithm can be run is 200kb. At this resolution, the window can be partitioned into two separate TADs of 5 bin width. However, this is inappropriate as previous research indicated that TADs do not begin truly appearing until the resolution becomes less than 100kb [11]. Therefore, our method is viable for all potential resolutions from which meaningful TADs can be called.

The algorithm starts at the beginning of the matrix and identifies the TADs in the first window. The window is then moved forward to the beginning of the last TAD detected, to account for the fact that the final TAD may overlap between windows. This is repeated until the end of the matrix. The result is a unique set of TADs.The first step of the algorithm is to find the graph spectrum. First, we calculate a Laplacian matrix - a matrix containing the spatial information of a graph. Multiple Laplacians exist [56]; but since our method builds upon the multiclass spectral clustering algorithm [33], which uses the symmetric Laplacian, we use the normalized symmetric Laplacian as follows:

1. Calculating the normalized symmetric Laplacian:

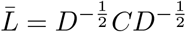

where *D* = *diag*(**1**^*T*^*C*)
2. Solve the generalized eigenvalue problem:

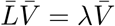 The result is a matrix of eigenvectors 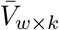, where *w* is the window size, and *k* is the number of eigenvectors used, and a vector of eigenvalues where each entry *λ_i_* corresponds to the *i_th_* eigenvalue of the normalized Laplacian 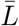.
3. Normalize rows and columns to sum to 1:

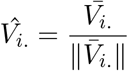

where the subscript *i*. corresponds to column *i*.

#### Projection onto the unit circle

Our method builds on the approach to spectral clustering first introduced in [33], which works by projecting the eigenvectors on a unit circle. Once we project these values on the circle, we can cluster regions of the genome by simply finding gaps in the circle (Supplemental Figure S9). In the unit circle representation, a TAD boundary can be thought of as a region of discontinuity in the eigenvectors of adjacent values. Regions within the same TAD should have similar eigenvectors and have small distances between them. This approach takes advantage of the fact that eigenvectors are mapped to genomic coordinates which have a natural ordering. The steps for this portion of the algorithm are below:

1. Normalize the eigenvectors and project onto a unit circle:

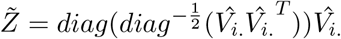
2. For *i* = 2, …, *n* where *n* is the number of rows in 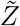 and *k* is the number of eigenvectors calculated (We suggest using two) to produce 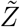, calculate the Euclidean distance vector *D*:

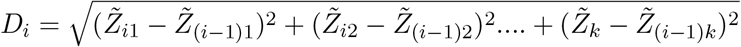 This step calculates the distance between the entries of the first two normalized eigenvectors that are associated with bin *i* and the bin to its left.

#### Choosing the number of TADs in each window

1. Find the location of the first 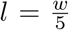 largest values in *D_i_*, where *w* is the window size, and *l* + 1 is the maximum number of TADs in a given window and partition the matrix into *l* + 1 sub-matrices with boundaries defined by the location of the *l* largest values.
2. For each sub-matrix calculate the silhouette statistic [35]:

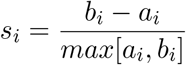 Here, *a* is the mean distance between each cluster entry and the nearest cluster and *b* is the mean distance between points in the cluster. The distance between two given loci *i* and *j* is defined as 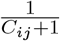, with “+1” added to avoid division by zero. *C_ij_* corresponds to the number of contacts between loci *i* and loci *j*.
3. Find the mean silhouette score over all possible numbers of clusters *m* and organize into a vector of means:

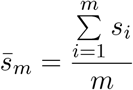
4. Find the value of *m* which maximizes 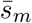

By taking the mean silhouette score, we can determine the number of eigenvectors which allows us to maximize the similarity within clusters while minimizing the similarity between clusters. This translates into the number of clusters (i.e., TADs) that produces the most well-separated clusters. This procedure is performed within each window, further allowing to identify poorly organized regions (gaps, Supplemental Methods). Cluster boundaries are mapped to genomic coordinates based on their location in the contact matrix. If a TAD is detected and found to be less than 5 bins wide it is ignored due to previous evidence suggesting these are not biologically relevant [11,53–55]. This step implies that, for a given window, the maximum number of TADs in a window is equal to the size of the window divided by 5.

#### Creating a hierarchy of TADs

We can find a hierarchy of TADs by iteratively partitioning the initial TADs. This is done by running a modified version of the main algorithm that includes an extra filtering step that tests for the presence of sub-TADs in each TAD. Briefly, each TAD is treated as an individual contact matrix, and a window is not used. To test for the existence of sub-TADs, we convert the distance vector *D_i_* into a set of Z-scores by first taking the natural log of the distance vector before centering and scaling. This is done following our empirical observation of the log-normality of eigenvector gaps (Supplemental Figure S10). We then label any distance with a Z-score greater than 2 as a sub-boundary. If significant sub-boundaries are detected, we partition the TAD with each sub-boundary indicating the end of a given sub-TAD. This procedure is then repeated for each sub-TAD until either the TAD is too small to be partitioned into two sub-TADs or if no significant boundaries are found. The TADs detected during the initial run of the algorithm are considered primary TADs and the TADs detected after partitioning are considered secondary, tertiary, etc., sub-TADs. In practice, this approach can also be used for the first iteration of the algorithm (non sub-TADs) and is an option in the `SpectralTAD` R package.

#### Benchmarking TAD callers

To evaluate the robustness of TAD callers simulates Hi-C matrices were systematically modified to contain pre-defined levels of noise, sparsity, and sequencing depth. The overlap of TAD boundaries with the gold-standard TADs was tested using Jaccard statistics; the cross-resolution TAD comparison was assessed using a modified version of the Jaccard coefficient (Supplemental Figure S1). The effect of Hi-C data normalization was tested using the iterative correction and eigenvector decomposition (ICE) [41], Knight-Ruiz (KR) [3, 42], and the Square Root Vanilla Coverage (sqrtVC) [3] methods. The association of TAD boundaries with genomic annotations, such as CTCF, was assessed using a permutation test. See Supplemental Methods for details.

## Software availability

The software is published under the MIT license. The source code of `SpectralTAD` is available at https://github.com/dozmorovlab/SpectralTAD.

## Acknowledgements

*Author contributions:* MGD and KGC conceived the project, KGC implemented `SpectralTAD` and wrote the analysis scripts. JCS helped with the data analysis. MGD and KGC wrote the manuscript with JCS contributions.

## Supplemental Legends

**Supplemental Figure S1. Example of modified Jaccard statistics to measure agreement between TAD boundaries detected at different resolutions.** The top triangles indicate TADs detected at 50kb resolution, while the bottom triangles indicate those detected at 25kb resolution. There are four shared boundaries (blue lines) and two non-shared boundaries (red lines). The traditional Jaccard statistic underestimates the fact that the four TAD boundaries agree at a different resolution, while the modified Jaccard statistics correctly identifies the perfect overlap between TAD boundaries by ignoring resolution differences.

**Supplemental Figure S2. Examples of TADs detected by different TAD callers.** TADs detected by SpectralTAD, TopDom, OnTAD, HiCSeg and rGMAP. Darker red colors indicate a higher level of connectivity while lighter blues indicate less connectivity; triangles indicate TADs. Data are shown from [3], resolution 50kb, chr22 (Coordinates 23650000:26750000). All parameters were set according to the instructions of each package for analyzing 50kb resolution data with no normalization performed.

**Supplemental Figure S3. The effect of data normalization on the average number (A) and width in kilobases (B) of TADs and the average number of peaks in enriched markers (C) & depleted markers (D), enrichment (F) and depletion (G) for different genomic annotations.** Counts (A) and widths (B) for raw, KR-, ICE- and sqrtVC-normalized Gm12878 data at 25kb and 50kb resolutions, averaged across chromosome 1-22, are shown for primary, secondary, and tertiary TADs detected by SpectralTAD. The average number of annotations for enriched (D) and depleted (E) genomic features and the permutation p-values corresponding to enrichment (F), and depletion (G) for the top most enriched/depleted genomic annotations (permutation test) at TAD boundaries for Gm12878 data at 50kb resolution are shown.

**Supplemental Figure S4. The comparison of SpectralTAD and other TAD callers regarding TAD consistency and biological significance.** To test for robustness to noise, sparsity, and downsampling, TADs were called from simulated Hi-C matrices using SpectralTAD and other TAD callers. The TAD boundaries were extended by 50kb regions flanking a boundary on both sides. They were compared with the ground-truth TADs using the Jaccard similarity metric. The performance of the TAD callers was assessed at different level of noise (A, the percentage of the original contact matrix modified by adding a constant of two), sparsity (B, the percentage of the original contact matrix replaced with zero), and downsampling (C, the fraction of contacts kept, see Methods). Using the raw data from Gm12878 at 50kb resolution, enrichment of genomic annotations within 50kb regions flanking a TAD boundary on both sides was assessed using a permutation test. The average number of annotations for enriched (D) and depleted (E) genomic features and the permutation p-values corresponding to enrichment (F), and depletion (G) for the top five most enriched/depleted genomic annotations are shown. Results averaged across chromosome 1-22 are shown.

**Supplemental Figure S5. The number, width, and consistency of TADs called across resolutions and primary vs. replicate for different methods.** The average number (A) and width (B) of TADs across resolutions, Jaccard similarity between TAD boundaries detected from primary and replicate data and modified Jaccard similarity between TAD boundaries detected from data at 10kb, 25kb and 50kb resolutions (C). Wilcoxon test p-values are shown. Data from Gm12878 cell line, chromosomes 1-22.

**Supplemental Figure S6. The effect of the hierarchy of TAD boundaries detected by SpectralTAD on the average number of annotations in enriched (A) & depleted (B) genomic markers and on enrichment (C) and depletion (D) for different genomic annotations.** Results for TAD boundaries detected as level 1, 2, and 3 boundaries are shown. Genomic annotations were considered within 50kb regions flanking a boundary on both sides. Wilcoxon test p-values are shown in panel A & B and aggregated p-values, using the Fisher’s method, are shown for panels C & D. Raw data from Gm12878 cell line, chromosome 1-22, 50kb resolution.

**Supplemental Figure S7. Jaccard similarity of TAD boundaries across cell types (A) and tissues (B).** TADs were called using SpectralTAD. Clustering was performed using Ward clustering applied to a Jaccard distance matrix. All TADs were called on raw 40kb data from [49]. Various cell-lines and tissues are used.

**Supplemental Figure S8. Runtime performance of various TAD callers.** TADs were called using data from Gm12878 cell line at 10kb and 25kb resolution, and runtimes recorded. (A) Runtimes were summarized across different chromosomes. Each dot represents chromosome-specific run time averaged across three runs, with the regression line approximating the trend. X-axis - chromosome size in the number of bins, Y-axis – time in seconds. (B) The total time to analyze chromosomes 1-22 was calculated and summarized across methods and levels of parallelization for Gm12878 25kb resolution data. X-axis - Method, Y-axis – time in seconds. Results for HiCSeg are excluded due to exceptionally slow runtimes (24+ hours for one 10kb chromosome).

**Supplemental Figure S9. Projection of eigenvectors on the unit circle.** This projection allows us to identify TADs based on the distance between eigenvectors. The two largest gaps are used to seperate TAD 1, TAD 2 and TAD 3. We can also see the difference between a strongly organized group with close together points (TAD 1 and TAD 2) and a weaker group with more spread out points (TAD 3). Simulated data from [50]

**Supplemental Figure S10. Distribution of eigenvector gaps.** The distributions of eigenvector gaps are plotted separately for each 10kb, 25kb and 50kb contact matrix from [3], 131 chromosome-specific datasets total. Results are colored by resolution. Higher resolution data shows smaller overall gaps due to a larger number of regions of high sparsity. The untransformed eigenvector gaps (A) and the natural log eigenvector gaps (B) are shown. MASS::fitdistr() function was used to establish the best fit by a lognormal (67 datasets) or a Weibull (64 datasets) distributions with similar log-likelihoods. The lognormal fit was chosen to model the distribution of log eigenvector gaps.

**Supplemental Table S1. Summary of gaps.** The percentage of gaps is summarized for all chromosomes at 10kb, 25kb, and 50kb resolution using raw GM12878 data [3]. Gaps are separated based on whether they are centromeric or other (unsequenced, or poorly organized chromatin).

**Supplemental Table S2. Experimental Data sources.** Genome annotation (hg19/GRCh37) [51] data for Gm12878 cell line used in the analysis, sorted by category, then by data type.

**Supplemental Table S3. Enrichment by Method.** Enrichment/Depletion results are provided for all genomic annotations tested. Permutation p-values summarized using Fisher’s method are shown. Data is sorted alphabetically by category and then by genomic annotation.

**Supplemental Table S4. Hi-C Data sources.** Information about experimental [3, 49] and simulated [50] Hi-C data.

**Supplemental Table S5. Jaccard similarity across TAD hierarchy.** Results for the corresponding comparison of Primary, Secondary, Tertiary TADs and Level 1, 2, 3 TAD boundaries are shown. Jaccard similarity coefficients were compared using a Wilcoxon signed rank test. Column p-values correspond to the comparison of Jaccard within levels between tissue samples and cell-lines. Row p-values correspond to the comparisons within each type of data across hierarchy.

