## Supplementary material for "SpectralTAD: an R package for defining a hierarchy of Topologically Associated Domains using spectral clustering": Supplemental_Table_S5.pdf

| | Primary vs. Secondary | | | Primary vs. Tertiary | | | Secondary vs. Tertiary | | |
|---|---|---|---|---|---|---|---|---|---|
| | Primary | Secondary | P-value | Primary | Tertiary | P-value | Secondary | Tertiary | P-value |
| Cell Line | 0.42 | 0.40 | 0.0001 | 0.42 | 0.35 | <0.0001 | 0.40 | 0.35 | <0.0001 |
| Tissue | 0.22 | 0.21 | 0.0003 | 0.22 | 0.18 | <0.0001 | 0.21 | 0.18 | <0.0001 |
| P-value | <0.0001 | <0.0001 | | <0.0001 | <0.0001 | | <0.0001 | <0.0001 | |

| | Level 1 vs. Level 2 | | | Level 1 vs. Level 3 | | | Level 2 vs. Level 3 | | |
|---|---|---|---|---|---|---|---|---|---|
| | Level 1 | Level 2 | P-value | Level 1 | Level 3 | P-value | Level 2 | Level 3 | P-value |
| Cell Line | 0.23 | 0.23 | 0.0347 | 0.23 | 0.30 | <0.0001 | 0.23 | 0.30 | <0.0001 |
| Tissue | 0.12 | 0.10 | <0.0001 | 0.12 | 0.13 | 0.0600 | 0.10 | 0.13 | <0.0001 |
| P-value | <0.0001 | <0.0001 | | <0.0001 | <0.0001 | | <0.0001 | <0.0001 | |
