## Supplementary material for "SpectralTAD: an R package for defining a hierarchy of Topologically Associated Domains using spectral clustering": Supplementary_Material.pdf

### Supplemental Material

#### Previous Methods

Many of the first and most popular TAD detection methods were based on the directionality index which is a function of the average upstream and downstream interactions. This metric was then used as a parameter in a hidden Markov model, to establish the location of TAD boundaries (Dixon et al. 2012). The basis of this method was the fact that boundary regions are expected to interact with downstream regions more than upstream regions. This method was followed by a number of methods designed to calculate non-hierarchical TADs such as `Armatus` (Filippova et al. 2014), `HiCseg` (Lévy-Leduc et al. 2014), `TADLib` (Wang et al. 2015), `TopDom` (Shin et al. 2016), `Arrowhead` (Durand et al. 2016), `TADbit` (Serra et al. 2017), `CaTCH` (Zhan et al. 2017) and `RHiCDB` (Chen et al. 2018). Another intuitive metric, the insulation index (Crane et al. 2015), uses a sliding window approach to sum up contacts within a given region surrounding each locus. As TADs are regions of increased contacts, they can easily be identified via a contact count cutoff. Some tools, such as the `TADtool` Python package (Kruse et al. 2016), implement both metrics to call TADs. `HiCDB` uses an extension of the conventional insulation index that corrects for background noise.

The detection of hierarchical TAD structures was first introduced by Fraser J. et al. (Fraser et al. 2015) who used single-linkage clustering to create a hierarchical structure of meta-TADs which contain smaller sub-TADs. Since this discovery, there has been a lack of publicly available, user-friendly, hierarchical TAD callers. A hierarchical TAD caller refers to a tool that finds TADs and sub-TADs contained within them. To date, the choice of hierarchical TAD callers remains limited.

The first publicly available tool for hierarchical TAD calling, `TADtree` (Weinreb and Raphael 2016) worked by creating TAD "forests" containing hierarchical "trees" of TADs. `TADtool` similarly provides hierarchical TAD detection and visualization (Kruse et al. 2016). Other hierarchical TAD finders include `HiTAD` (Wang et al. 2017a) and `IC-Finder` (Haddad et al. 2017) which take dynamic programming, hidden Markov model-dynamic programming hybrid, and probabilistic approach, respectively. `ClusterTAD` (Oluwadare and Cheng 2017) introduced a traditional hierarchical clustering-based approach to TAD classification. Another method, `rGMAP` (Yu et al. 2017) has arisen as a potentially useful tool for TAD detection. This method utilizes a Gaussian Mixture model and a z-test of proportions to identify TADs. The model is then run iteratively to partition TADs into sub-TADs, but in practice is limited to two levels of TADs. More recently, a method called `onTAD` was proposed (An et al. 2018). This method uses `TopDom`, a single-level TAD caller that uses a statistical test on upstream and downstream contacts, to find all possible TAD boundaries and then to select a final configuration using a dynamic programming algorithm (Shin et al. 2016). Of these methods `TADtool`, `TADtree` and `TADLib` are Python based. `IC-Finder` and `ClusterTAD` are MATLAB based with `ClusterTAD` also including Java implementation and rGMAP is available as an R package. Although some comparison of TAD detection tools has been performed (Ay and Noble 2015; Forcato et al. 2017; Nicoletti et al. 2018), this diversity leaves the choice of the most appropriate method uncertain.

Hi-C data, represented as an adjacency matrix, naturally lends itself to the use of graph theory (Boulos et al. 2013; Wang et al. 2017b, 2013). `Arboretum-HiC` first introduced the idea of using Laplacian-graph segmentation to find structures in Hi-C data (Fotuhi Siahpirani et al. 2016). This method used a spectral clustering approach to simultaneously find 3D structures between multiple matrices. Chen et al. (Chen et al. 2016) proposed a method that framed the contact matrix as a weighted adjacency matrix and used recursive partitioning of the Fiedler vector to identify TADs. `MrTADFinder` (Yan et al. 2017) and `HiTAD` (Wang et al. 2017a) take a similar approach but address the question as a community detection problem. Most recently, `3DNetMod` was introduced which treats the Hi-C matrix as a network and uses network modularity to cluster the TADs (Norton et al. 2018). This method is also designed to find hierarchies of TADs. Network theoretical, or more broadly graph theoretical, approaches have a promise to provide us with a data-driven method of identifying TADs that takes the entire structure of loci-loci interactions into account.

R/Bioconductor is the de facto gold standard programming language for the genomics community (Gentleman et al. 2006). Currently, the number of TAD callers implemented in the R programming language are limited and include `HiCseg` (Lévy-Leduc et al. 2014), `TopDom` (Shin et al. 2016), `rGMAP`(Yu et al. 2017), and `HiCDB` (Chen et al. 2018). `HiCseg` is the only TAD-calling specific R package available on CRAN or Bioconductor, while `TopDom` is a downloadable R script. `HiCDB` and `rGMAP` are available on GitHub. Another tool, `RobustTad`, is in development and currently only provides a metric for TAD calling. It will potentially provide a new option for R users (Dali et al. 2018). Additionally, the `HiTC` R package (Servant et al. 2012) has functionality for calculating the directionality index but does not provide any tools for TAD identification. Neither `TopDom` nor `HiTC` can operate on the commonly used $n \times n$ contact matrices in text format. `TopDom` requires the data to be formatted as an $n \times n + 3$ matrix with the first three columns corresponding to the genomic coordinates. `HiTC` requires the user to transform the data into their package-specific `HTCexp` object. `HiCseg` can be used to analyze $n \times n$ text matrices but forces users to assume a distribution of contacts and estimate the number of TADs before running. These factors aren't always clearly apparent from the data and thus require constant tweaking to account for different levels of noise, sparsity, and resolution of Hi-C data. Although `HiCDB` can process $n \times n$ contact matrices, it requires matrices from multiple chromosomes thus limiting the analysis of single-chromosome data. Additionally, it requires data to have a resolution of 5kb, 10kb or 40kb. The aforementioned limitations limit the choice of R-based tools for direct comparison.

#### Identification and removal of gaps

We define gaps as regions where there is no coherent connectivity structure. There are two types of gaps, those with no contacts at all (centromeres and unsequenced regions), and regions where contacts exist but TADs are not present. The first category of gaps is removed by simply getting rid of loci with more than 95% zero contacts. Since there are situations when TADs span unsequenced regions, we allow TADs to start on one side of a gap and end on another. This is done by essentially treating regions on either side of a gap as being adjacent. The second category of gaps is detected using the silhouette score. Regions where contacts are present but no TADs exist will frequently have low silhouette scores due to the poor similarity between each locus within the region. As a result, we can detect this type of gap by treating it as a potential TAD then filter it out if its silhouette score is lower than .25.

#### Simulating levels of noise, sparsity, and sequencing depth

Simulated contact matrices (Forcato et al. 2017) (supplemental Table S4) were modified to simulate various levels of noise, sparsity, and sequencing depth. The noise was added by randomly selecting a percentage (4%, 8%, 12%, 16%, and 20%) of entries in the matrix and adding a constant of two to these entries. Entries were sampled with replacement meaning certain entries may have received more noise than others. In summary, we used five replicates at each noise level, totaling 25 simulated matrices.

We created two extra sets of contact matrices for simulating sparsity and sequencing depth. To mimic sparsity, we took the five simulated matrices with the minimum level of noise (4%) and introduced 90%, 75%, 50%, 25%, or 10% of zeros uniformly at random, totaling 25 matrices. To simulate changes in sequencing depth, we took the same five minimum noise matrices and applied the downsampling procedure adapted from (Yardimci et al. 2017). Briefly, the full contact matrix was converted into a vector of pairwise individual intra-chromosomal contact counts. The vector was downsampled uniformly at random proportional to the level of downsampling (1/2, 1/4, 1/8, and 1/16). Following downsampling, the vector was re-binned to the original contact matrix. This procedure produced a set of 20 matrices (five matrices, each modified by four levels of downsampling) with varying levels of downsampling.

#### Normalization of Hi-C data

Normalization of Hi-C data matrices is a common step in Hi-C data analysis (Lieberman-Aiden et al. 2009; Imakaev et al. 2012; Knight and Ruiz 2012; Cournac et al. 2012; Hu et al. 2012; Li et al. 2015; Ay et al. 2014). To test for the effect of normalization on the detection of TADs, we applied iterative correction and eigenvector decomposition (ICE) (Imakaev et al. 2012), Knight-Ruiz (KR) (Rao et al. 2014; Knight and Ruiz 2012) normalization, and the Sequential Component Normalization (SCN) (Cournac et al. 2012) to the simulated and real Hi-C matrices. ICE was implemented using the `ICE` function from the `dryHiC` R package version 0.0.0.9000. KR-normalization (Vidal et al. 2018) was performed using Juicer (Durand et al. 2016), and SCN normalization was implemented using HiCcompare version 1.5.0 (Stansfield et al. 2018).

#### Measuring association of TAD boundaries with genomic annotations

To test for the enrichment of a genomic annotation at a given TAD boundary, we measure the total number of annotations within 50kb of the boundary point on either side. The flanking region accounts for the fact that a TAD boundary is a single point. Flanking also helps to correct for impreciseness due to issues like overlapping TAD boundaries and differences in resolutions. We specifically choose 50kb for comparison of genomic features because it allows for at least one bin of wiggle room for all resolutions used in this paper. By keeping the size of the flanking region consistent across resolutions, we can directly compare enrichment at TAD boundaries across resolutions.

A permutation test was used to quantify the enrichment. Briefly, the mean number of genomic annotations within 50kb of each boundary was calculated for the entire chromosome. Two sets of bins, one the size of the TAD boundaries and another the size of all other regions were sampled without replacement, and the difference in the mean number of genomic annotations in the corresponding sets was calculated. This procedure was repeated 10000 times. We determined the permutation p-value by taking the number of randomly sampled (expected) mean differences that were greater than the observed difference in means between TAD boundaries and all other regions of the chromosome and dividing by 10000. $\alpha = 0.05$ was set to assess statistical significance. Given the fact that `TopDom` detects all TADs in one run while `rGMAP` and `SpectralTAD` detect hierarchical TADs, hierarchial TAD boundaries were combined for `rGMAP` (two levels) and `SpectralTAD` (three levels).

#### Jaccard and its modified version as a measure of similarity between TADs

Traditionally, Jaccard is used as a measure of overlap between sets. We use it as a measure of overlap between TAD boundaries. Given a set of TAD boundaries $A$ and $B$, we define the Jaccard as:

$$J = \frac{A \cap B}{A \cup B}$$

In plain terms, this is the set of shared boundaries divided by the total number of unique boundaries.

While we expect TADs called at different resolutions of the same Hi-C data to be nearly identical, TADs called from a higher-resolution data may be finer partitioned than those called from lower-resolution data. The traditional Jaccard measure will penalize for these finer TADs even though the original boundaries were detected. We introduce a modified Jaccard score of $J_a$, which accounts for the difference between TADs detected across resolutions.

$$J_a = \frac{A \cap B}{min(|A|, |B|)}$$

Here, $A$ and $B$ are two sets of TAD boundaries and $|A|$ and $|B|$ indicates the size of sets (supplemental Figure S1). This method is identical to the Jaccard statistic, but instead of dividing by the union of $A$ and $B$ we divide by the smallest size. A score of 1 indicates that all of set $A$ is contained in a subset of $B$ or vice-versa. This is in contrast to traditional Jaccard where a score of 1 indicates that all boundaries in $A$ and $B$ are identical.

##### Modified Jaccard with a flank

The modified Jaccard can be extended to account for the impreciseness of TAD callers or differences in the resolution which make finding identical TADs impossible. For instance, half of the loci in a 25kb resolution contact matrix aren't actually in the 50kb resolution version of the same matrix but in one of the neighboring bins. This "off-by-one" error is accounted for by extending TAD boundary points at higher resolution by flanking regions of size $f$. Consequently, the modified Jaccard formula becomes:

$$J_a = \frac{\mathbf{A} \cap \mathbf{B}}{min(|A|, |B|)}$$

where $\mathbf{A} = \{A, A + f, A - f\}$ and $\mathbf{B} = \{B, B + f, B - f\}$.

When comparing the 50kb and 25kb resolution matrices, we can set $f = 25000$ to make up for any difference in resolution. Note that modified Jaccard is used only when comparing boundaries between different resolutions; otherwise, the traditional Jaccard statistic is used.

##### Runtime analysis

One general drawback of spectral clustering is the fact that it scales poorly to large matrices. Traditionally, the three main bottlenecks are 1) the creation of the distance matrix, 2) the creation of the Laplacian matrix, and 3) the eigenvalue decomposition. Letting $n$ equal the number of loci in the genome, the distance matrix creation has a complexity of $\mathrm{O}(n^3)$. The Laplacian matrix creation involves two matrix multiplication steps. The traditional multiplication step costs $\mathrm{O}(2n^3)$, and the eigendecomposition step costs $\mathrm{O}(n^3)$. In total, these bottlenecks result in computational complexity of $\mathrm{O}(4n^3)$ or more simply $\mathrm{O}(n^3)$, cubic complexity.

Our method manages to avoid all of these bottlenecks. The first bottleneck is avoided because the contact matrix is itself an adjacency matrix and doesn't require transformation to a distance matrix. The second bottleneck is solved by calculating relatively small Laplacian matrices for each window instead of calculating the Laplacian matrix of the entire contact map. The total number of windows for a given chromosome of size $n$ with window size $w$ is equal to $\sim \frac{n}{w}$ with some discrepancy for rounding and variable TAD size. Accordingly, the computational complexity of Laplacian matrix creation over a given number of windows is equal to $O(w^3 \frac{n}{w})$ or more simply $O(n)$. The third bottleneck is similarly addressed by the windowed approach. The window makes the computation complexity of the eigendecomposition equal to $O(Kw^2nm)$, where $K$ is the number of eigenvalues and $m$ is the number of iterations for convergence of the eigensolver. This reduces to $O(n)$. As a result of these steps, the windowed spectral clustering algorithm reduces to the computational complexity of $O(2n)$, or simply linear complexity $O(n)$.
