## Supplementary figures and images for "SpectralTAD: an R package for defining a hierarchy of Topologically Associated Domains using spectral clustering"

### Supplemental_Fig_S1.png

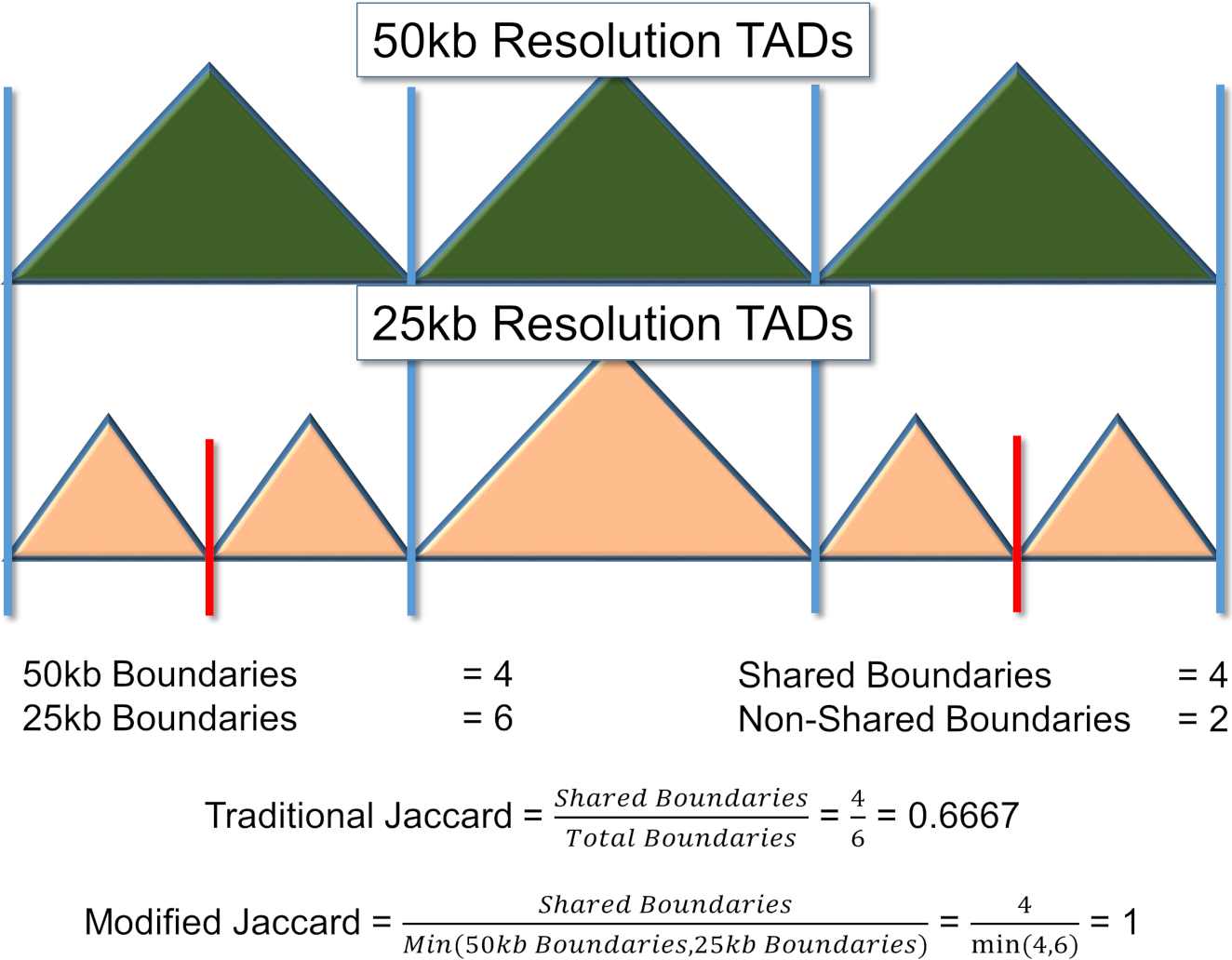

### Supplemental_Fig_S2.png

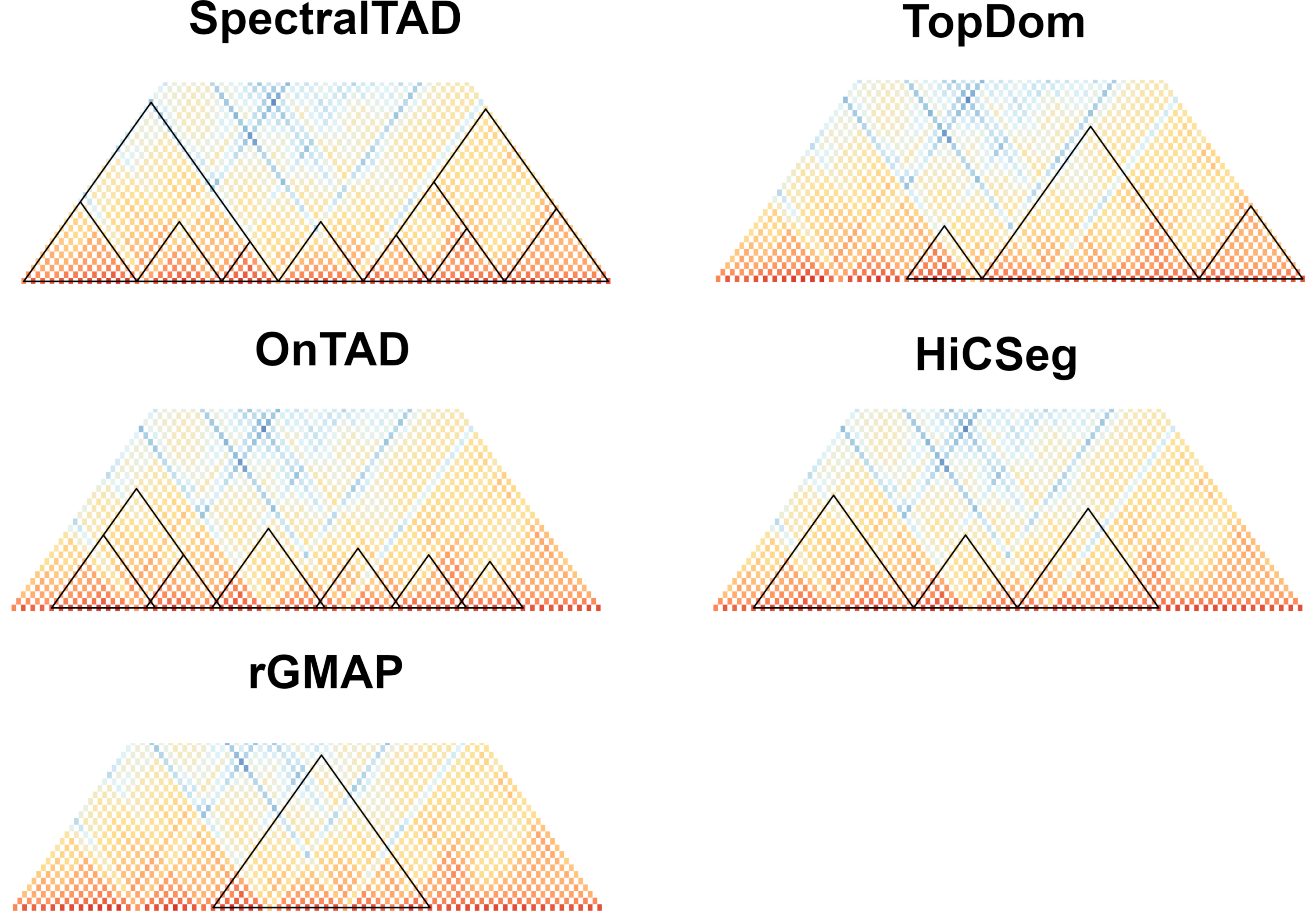

### Supplemental_Fig_S3.png

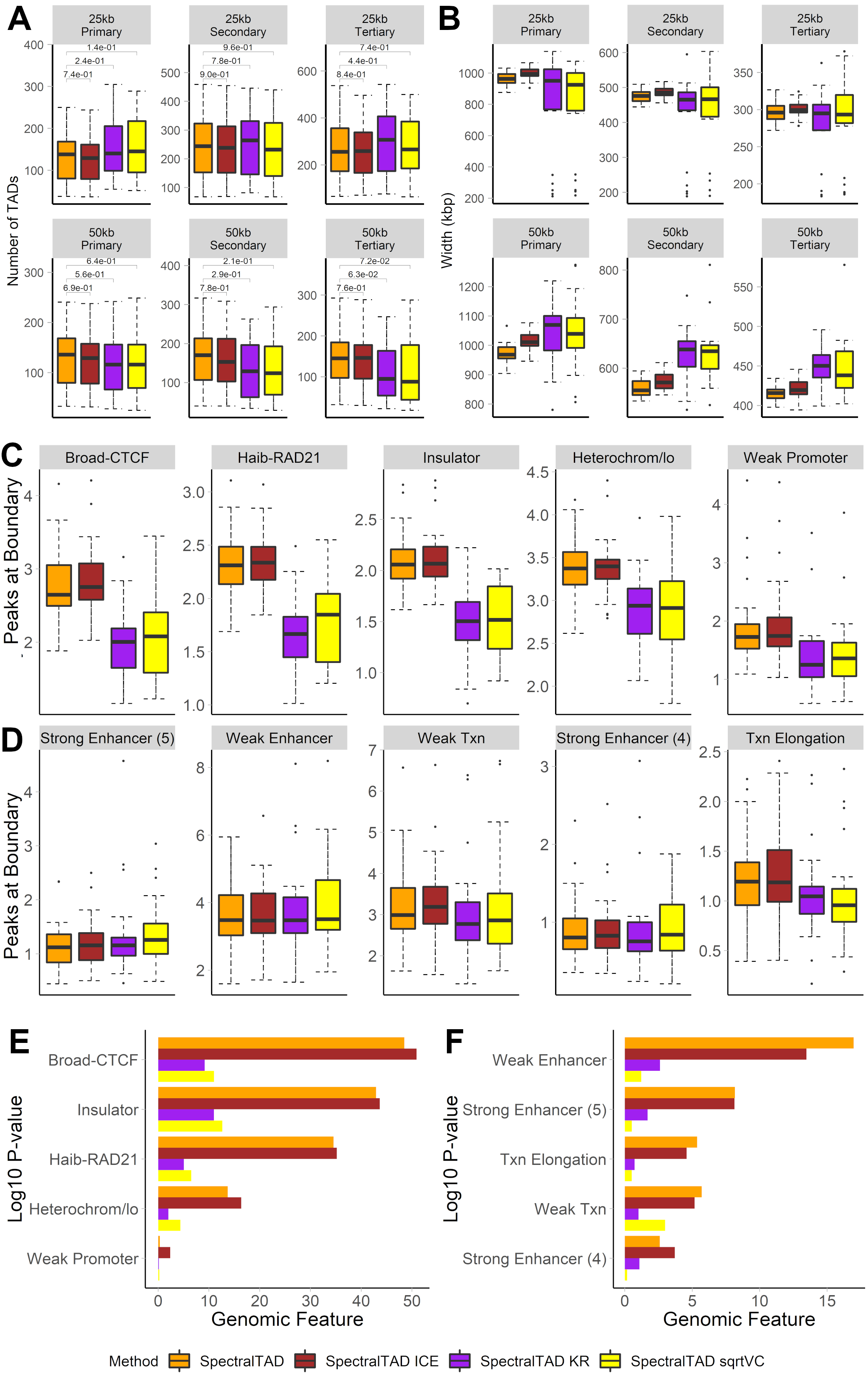

### Supplemental_Fig_S4.png

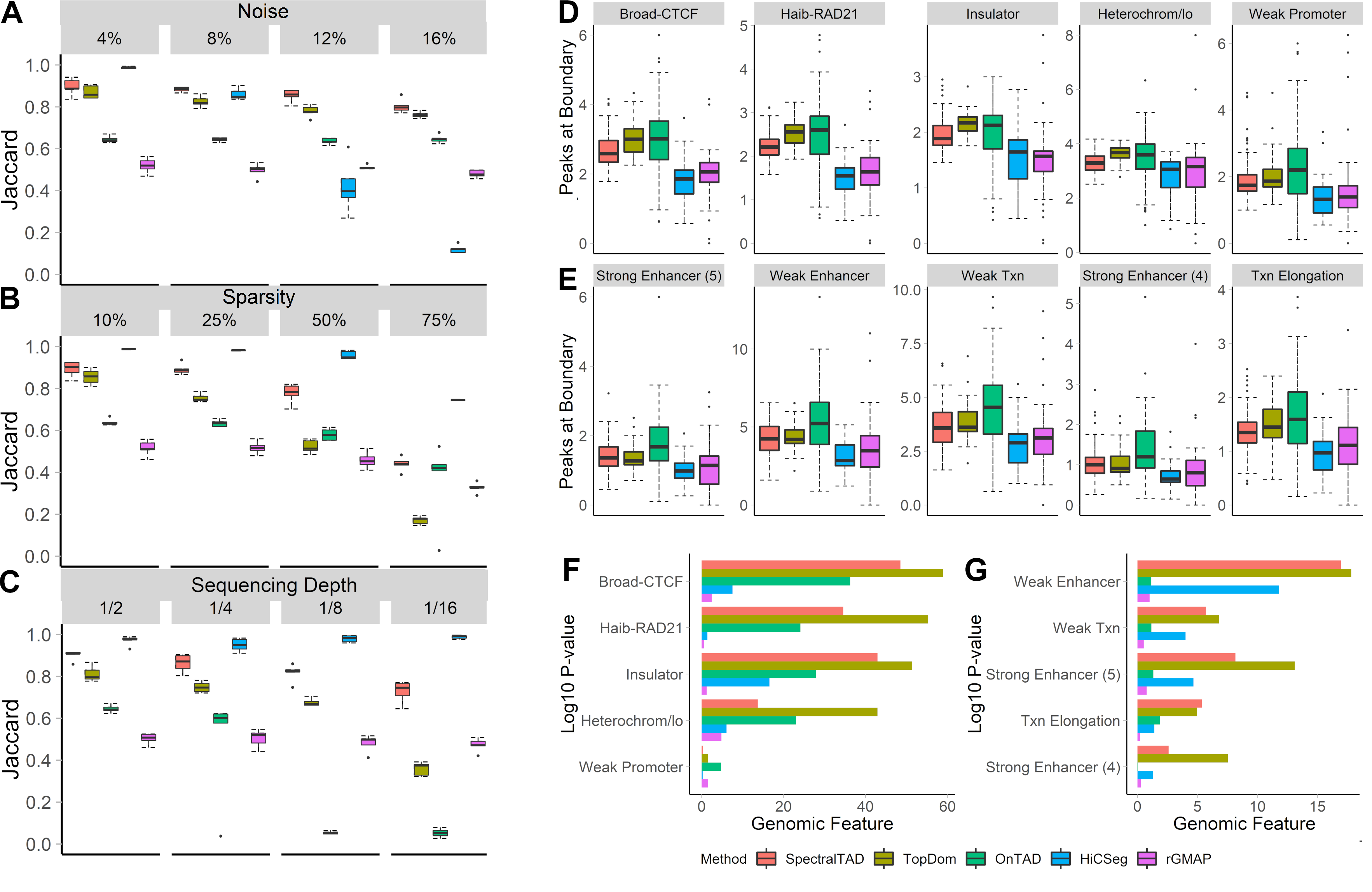

### Supplemental_Fig_S5.png

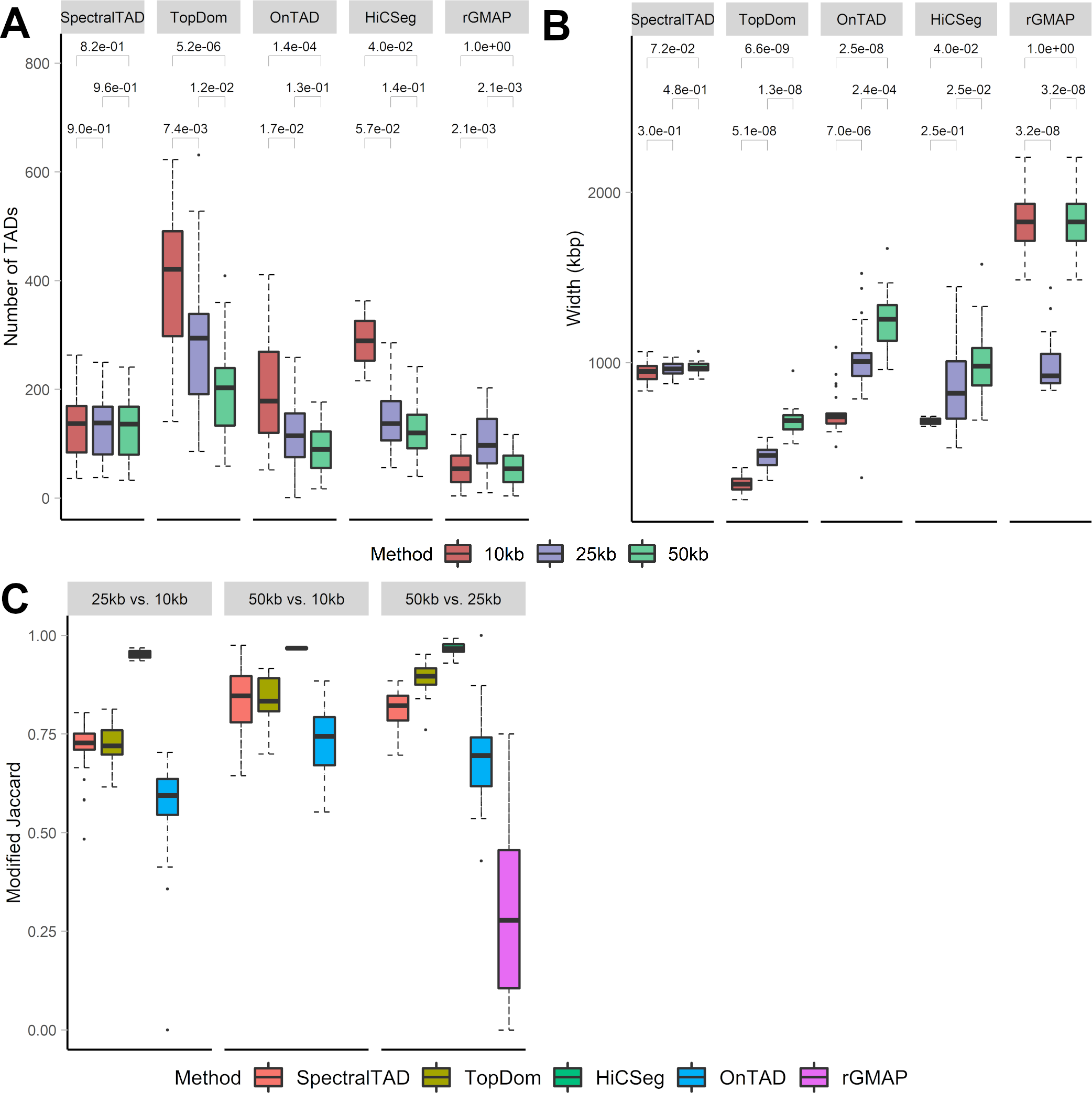

### Supplemental_Fig_S6.png

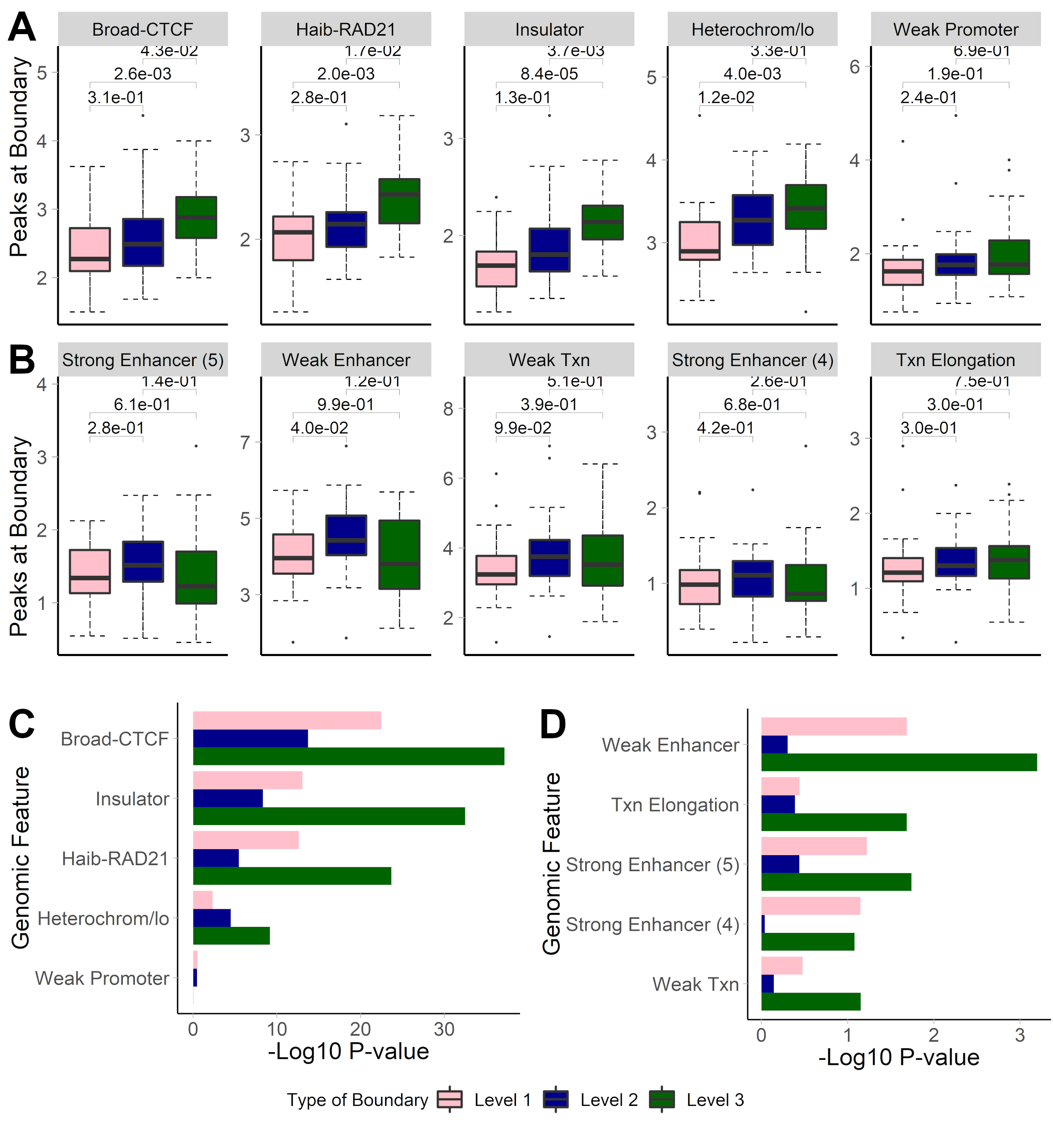

### Supplemental_Fig_S7.png

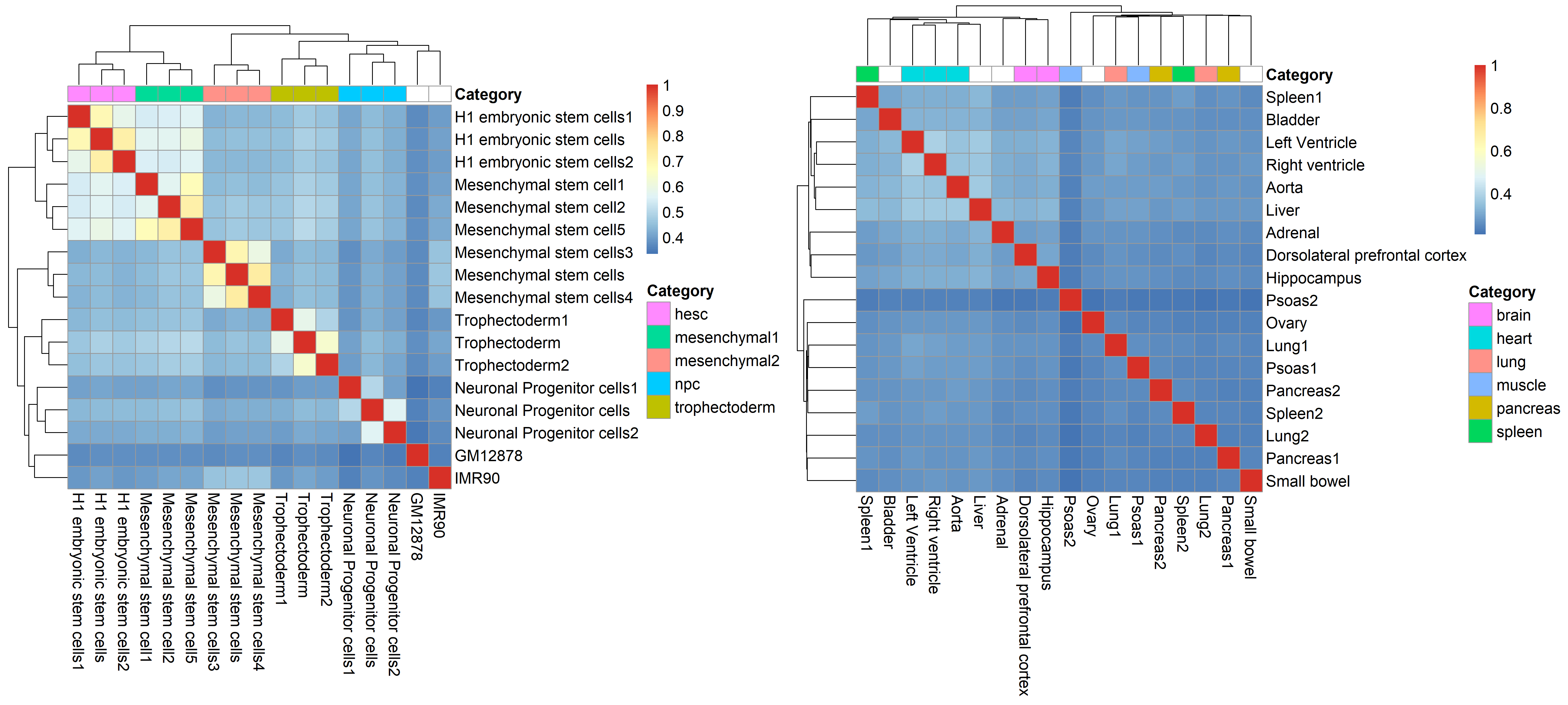

### Supplemental_Fig_S8.png

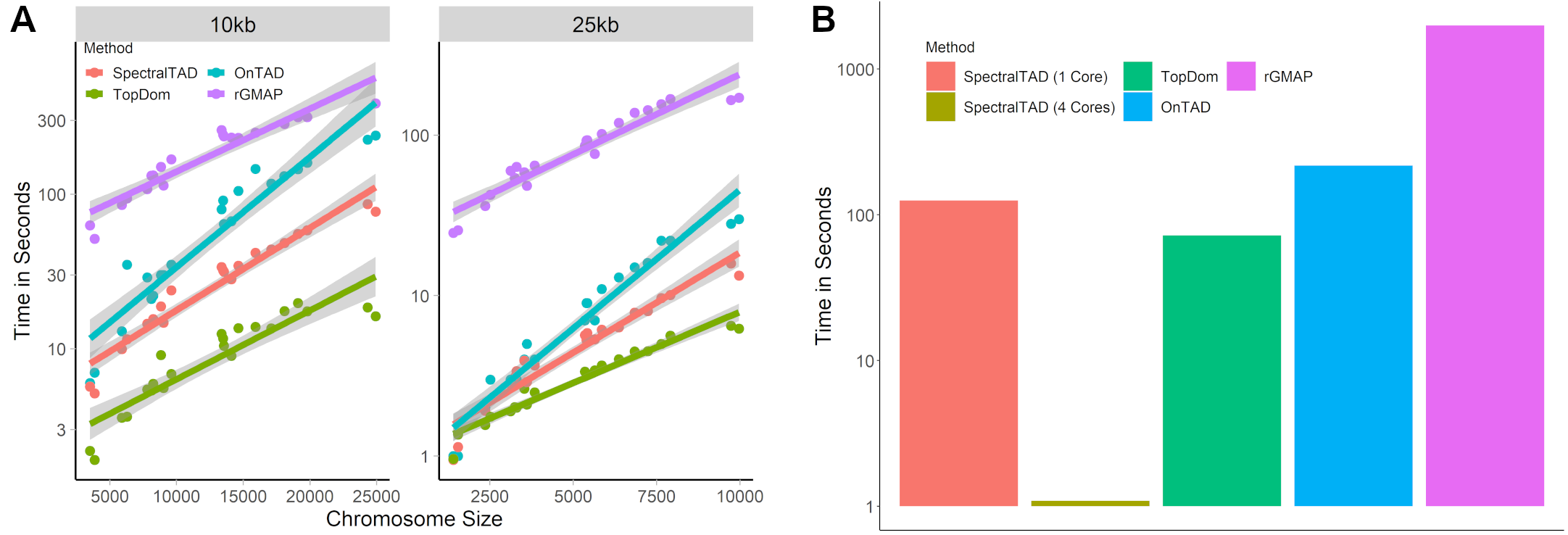

### Supplemental_Fig_S9.png

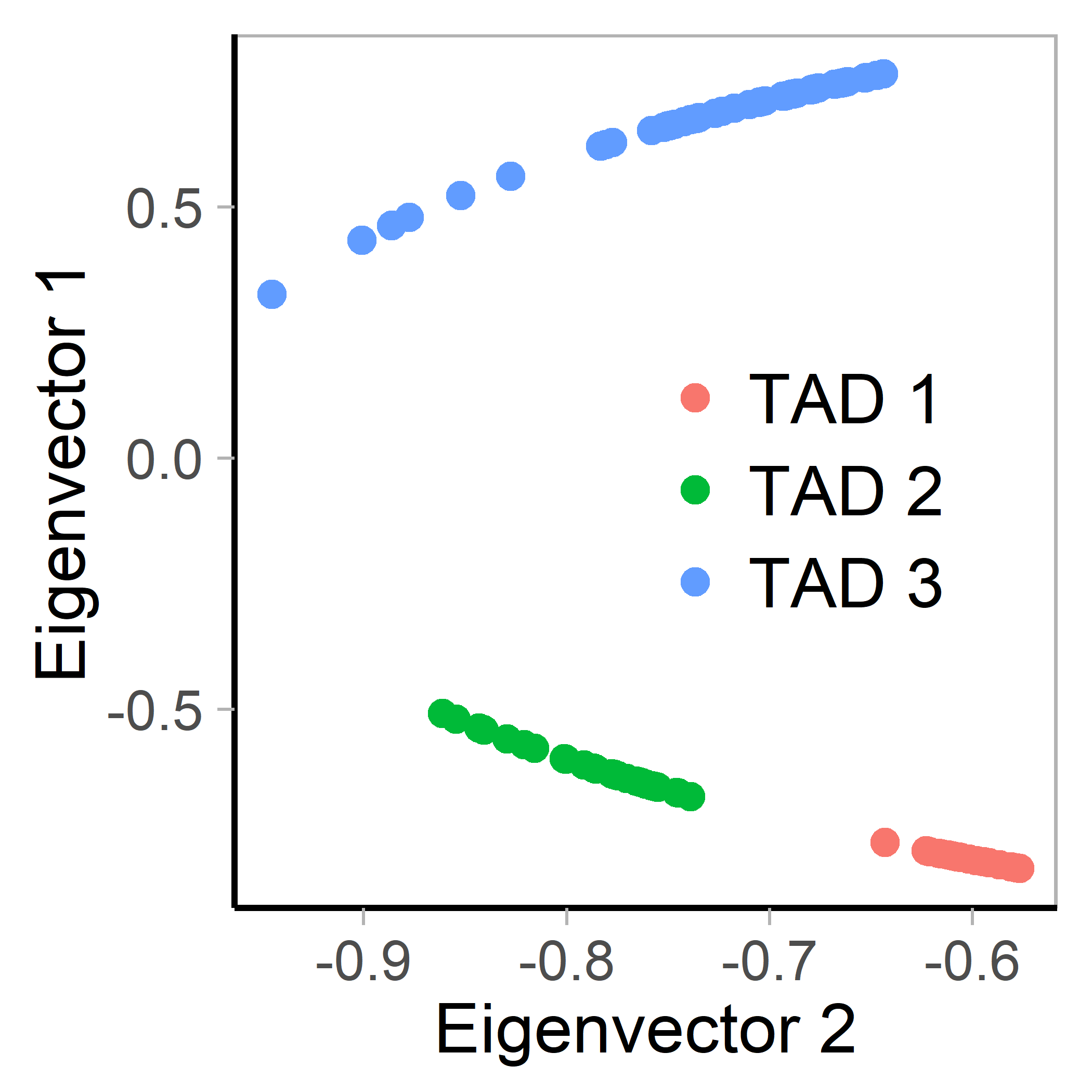

### Supplemental_Fig_S10.png

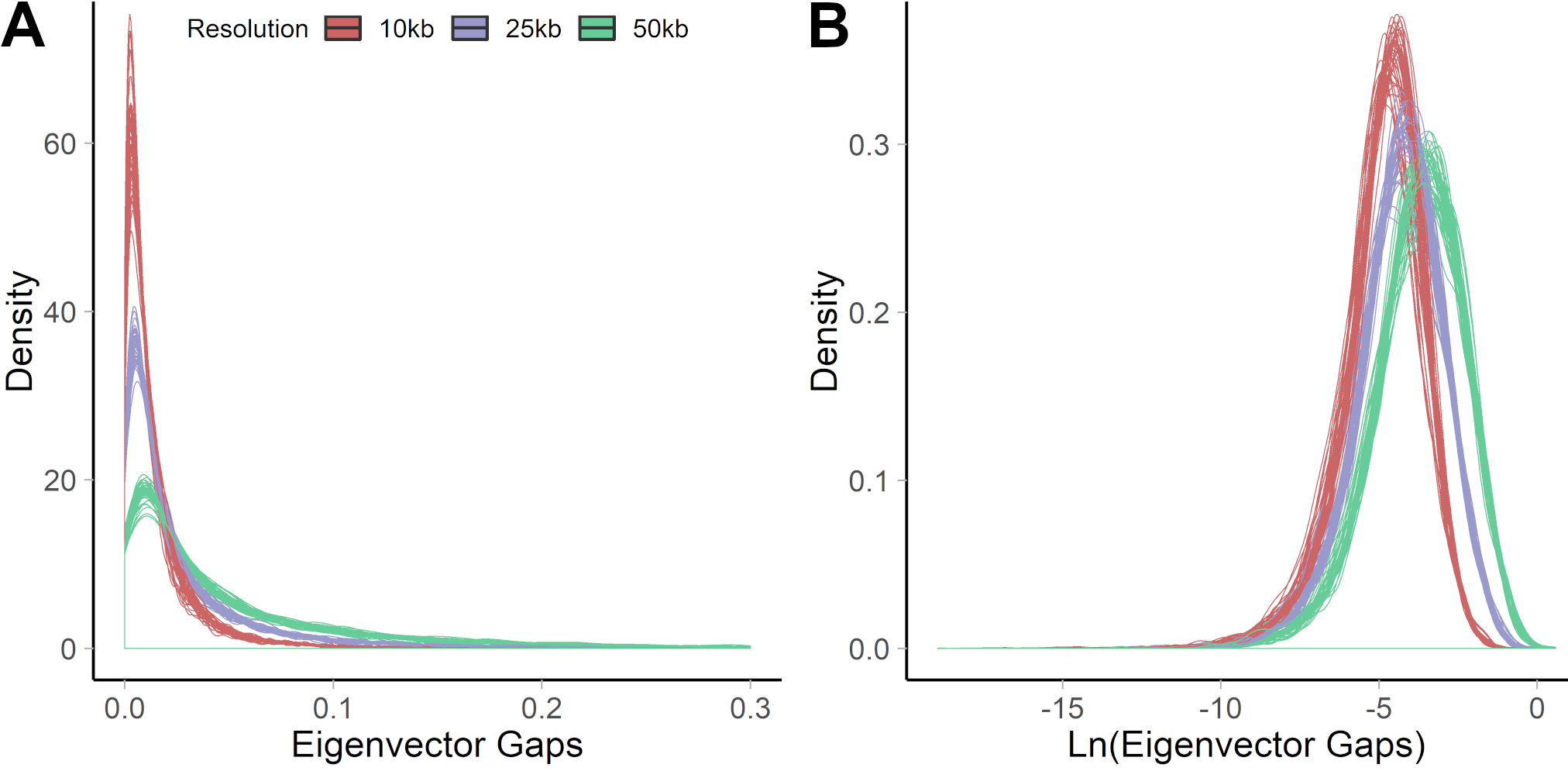
